# Multilevel HierCC typing scheme for global genomic epidemiology and antimicrobial resistance surveillance of *Campylobacter jejuni*

**DOI:** 10.64898/2025.12.17.694998

**Authors:** Ruochen Wu, Sandeep Kaur, Michael Payne, Li Zhang, Ruiting Lan

## Abstract

*Campylobacter jejuni* is the most common bacterial cause of human gastroenteritis around the world. The disease burden and financial cost associated with *C. jejuni* pose significant challenges to the global public health system. A stable, scalable and standardised typing scheme is essential for epidemiological and AMR surveillance for prevention and control of *C. jejuni* infections. We curated and assembled a *C. jejuni* global whole genome sequencing (WGS) dataset of 63,012 publicly available Illumina short-read genomes. A *C. jejuni* global cgMLST scheme with 1161 core loci was developed using a representative subset of 2,587 genomes. HierCC clustering of the cgMLST profiles was performed for all levels and suitable HierCC levels were selected to enable epidemiological analysis. Antimicrobial resistance (AMR) prediction was performed using AbritAMR. A HierCC cluster was classified as a resistant cluster if ≥80% of the isolates were resistant to at least one antibiotic.We selected 9 HierCC levels (HC2, HC4, HC6, HC12, HC19, HC55, HC100, HC167 and HC264) to form the multilevel HierCC scheme. The multilevel design allowed natural and genetically discrete clusters to be described consistently at both global and local scale. We showcased the utility of the scheme with identification of country and continent specific clusters at HC264 to HC19 levels and differentiation of animal host specific (specialist) from human infection (generalist) clusters at HC264 to HC55 levels. In the USA dataset from 2016 to 2022, a total of 276 resistant clusters were identified with 22 resistant clusters increasing in frequence in recent years. An increased prevalence of *C. jejuni* carrying *50S_L22* mutations in the USA was observed from 2016 to 2022, suggesting continuing selection pressure from macrolide use in the farm animals. We developed a novel multilevel HierCC typing scheme to standardise AMR surveillance and provide flexible typing resolution for both short- and long-term epidemiological investigations of *C. jejuni* at global and local levels.

## Introduction

*Campylobacter jejuni* is a motile, microaerophilic, Gram-negative bacterium with global prevalence. [1]. It is a zoonotic pathogen commonly recovered in most livestock and animal food products [2,3]. Human transmission predominately occurs through the consumption of undercooked or contaminated poultry products or unpasteurized dairy [3,4]. It is the leading bacterial cause of gastroenteritis in the world, impacting 96 million people annually by the estimation of world health organization (WHO) [5–7]. *C. jejuni* induced gastroenteritis has also been associated with long-term illness such as dysbiosis, irritable bowel syndrome (IBS), and Guillain-Barré syndrome (GBS) [8,9].

Antibiotics are rarely prescribed for *C. jejuni* infections as most cases are mild and self-limiting [4]. However, antibiotics are warranted for patients with severe gastrointestinal infections, with persistent fever, bloody diarrhea for seven days or more, significant volume loss and having eight or more bowl movements per day [4,10,11]. In such cases, the drug of choice is macrolides, specifically erythromycin or azithromycin, as recommended by American Academy of Paediatrics and Infectious Diseases Society of America [4,7,10–12].

In 2017, the EU issued an action plan to reinforce the need for comprehensive data collection, monitoring and surveillance of AMR [13]. The increasing emergence of antimicrobial resistance (AMR) and multi-drug resistance (MDR) in *C. jejuni* has been extensively documented, which has become a major public health concern in both developed and developing countries in recent years [7,14–17]. Antibiotic stewardship in the UK has been associated with some reduction in antibiotic resistance in farm animals, but the regulatory oversight in the USA has led to antibiotic overuse [18–22]. The impact of inadequate AMR surveillance and regulation on public health is likely to be more severe in developing countries [23].

Rapid advancement in next-generation sequencing (NGS) technologies has made whole genomes sequence (WGS) data more affordable and accessible than ever before [24]. Currently, the USA and the UK are the only two countries [25–27] that provided large amount of WGS data in the public domain from their national pathogen surveillance programs utilising NGS technology. Molecular typing techniques such as multilocus sequence typing (MLST) and core genome MLST (cgMLST) have contributed significantly to short- and long-term *C. jejuni* epidemiological studies respectively in recent years [28,29]. MLST has played a pivotal role in elucidating the genetic diversities of *C. jejuni* in both evolutionary and ecological studies [30,31]. The *C. jejuni* global population consisted of both globally and locally distributed MLST STs, with ST-21, ST45, and ST-50 being the most frequently sampled variant around the world [32,33]. The subsequent introduction of the Oxford cgMLST scheme for *C. jejuni* and *C. coli* has provided high resolution typing necessary for investigating local *C. jejuni* epidemiology [29,34–36]. However, both MLST and cgMLST schemes only offer fixed resolution, with no scalability. MLST lacks the resolution needed to differentiate closely related isolates, while the resolution of cgMLST alone could be too high for surveillance over longer time frames [37,38]

HierCC is a method that clusters cgMLST allelic profiles into multilevel hierarchical clusters [39]. Since 2018, HierCC has been implemented in EnteroBase to support a variety of functions across multiple hierarchical levels, from differentiating species/subspecies at low resolution levels to identifying individual transmission chains at high resolution levels [40]. The resolution flexibility of HierCC enables utility in both long-term and short-term epidemiological analysis, as demonstrated in *Salmonella, Escherichia/Shigella, Clostridioides, Vibrio, Yersinia*, and *Mycobacterium* in the EnteroBase [41]. HierCC generates thousands of clusters at every allelic difference level and there has been no simple means to determine which levels are most suitable for describing bacterial population structure and epidemiology. Therefore, here is a need for guidance or standardised recommendations to use a subset of the allelic levels for surveillance applications, which would allow standardised communications and comparisons across time and countries. In this study, we develop a cgMLST specifically for *C. jejuni* and identify nine HierCC levels to create a stable multilevel HierCC typing scheme that allows for scalable resolution with epidemiologically relevant divisions of the *C. jejuni* population. We defined country- or continent-specific clusters, farm animal-specific clusters, and resistant clusters to medically important antibiotics for the global *C. jejuni* population. Importantly, we demonstrated the utility of the multilevel HierCC typing scheme for AMR surveillance when incorporated with national WGS surveillance data from the UK and the USA.

## Methods

### Curation of Illumina genomic dataset

Publicly available *C. jejuni* raw Illumina paired end reads (n = 68,657) and metadata were downloaded and processed from the European Nucleotide Archive (ENA https://www.ebi.ac.uk/ena, June 2023) [42]. The processing and assembly of raw reads were performed using the shovill (v1.1.0) and strategic k-mer extension for scrupulous assemblies (SKESA) [43–48]. The assembled genomes were quality filtered based on thresholds in supplementary method 1.1. *Campylobacter* species confirmation of the assembled genomes was performed using the CampyGStyper [49].

### Development of the *C. jejuni* global cgMLST scheme

We developed a *C. jejuni* specific cgMLST using a global representative dataset to capture the global genetic variation of *C. jejuni.* Assembled genomes in the global dataset were assigned the pubMLST *Campylobacter* seven gene MLST STs using the MLST package (https://github.com/tseemann/mlst) [50]. The representative dataset was populated by randomly selecting one genome from each unique MLST STs (n=2,587) in the global dataset (n=63,102). The cgMLST scheme was constructed using the representative dataset and the Comprehensive and Highly Efficient Workflow BSR-Based Allele Calling Algorithm (chewBBACA v3.3.2) with default parameters. The core loci were further assessed for their typeability using the representative dataset (n=2,587). Only loci present in 95% of genomes in the representative dataset were retained in the scheme (detailed in Supplementary method 1.2). We named the final scheme the UNSW *C. jejuni* cgMLST scheme (available at: https://chewbbaca.online ).

### Implementation of HierCC in *C. jejuni* global dataset

HierCC v1.27 was used to cluster 63102 *C. jejuni* genomes based on the cgMLST allelic profiles at default setting [39]. The HierCC package HCCeval was modified in python to output the allelic profile distance matrix to reduce overall run time of large dataset. At each HierCC level, the silhouette score of the HierCC clusters and the distance matrix was calculated using scikit-learn package silhouette_score [51]. The adjusted rand score between the clustering of isolates at each HierCC level and the division of isolates in the seven gene MLST were calculated using scikit-learn package adjusted_rand_score [51].

### Antimicrobial resistance annotation

The genetic determinants conferring antibiotics resistance of each assembled genomes were identified using AbritAMR (version 1.0.14) at default setting [52]. The resistance determinant genes or mutations responsible for phenotypic resistance were only included with a coverage ≥ 90% and identities of ≥ 90%. Phenotypic interpretation of the 15 antibiotic classes and antibiotic combinations were predicted based on the mechanism of the resistance determinants. The validity of phenotype resistance prediction by AbritAMR was tested using WGS of 3,181 isolates with known phenotype AMR profiles from NCBI Pathogen Detection database in June 2023 (https://www.ncbi.nlm.nih.gov/pathogens). For macrolide resistance, only strains carrying *erm*(B) and mutations in the 23S rRNA gene are classified as resistant. Although mutations in *50S_L22* are also detected and annotated using AbritAMR, they are analysed separately as indicator of macrolides selection pressure, as they do no they do not confer high-level resistance by themselves.

### Resistance score, and its statistical comparison between different data subsets

The calculation of resistance score (RS) and statistical comparison of resistance rate of antibiotic classes or combinations was performed using python (version 3.9.6) script from AMR-analysis [53]. The RS value for each genome was calculated by converting the susceptibility to each of the 15 antibiotic classes into Boolean values, where susceptible was registered as 0 and resistance was registered as 1. The distribution of RS between different time frames were compared using the Mann-Whitney *U* test and the significance tested at a Bonferroni corrected *p*-value [54,55]. For each genome, a resistance pattern was calculated using the Booleanised representation of resistance in the 15 antibiotics. Thus, the pattern of 1,1,1,1,1,1,1,1,1,1,1,1,1,1,1 indicated resistance to all antibiotics and the pattern of 0,0,0,0,0,0,0,0,0,0,0,0,0,0,0 indicated susceptible to all antibiotic classes above. The antibiotics are ordered based on the sequence in supplementary method 1.3.

### Identification of resistant HierCC clusters

For each HierCC level, we calculated the percentage of resistant strains in each cluster. Resistant HierCC clusters were identified when over 80% of the isolates are predicted resistant to a given antibiotic. From the lowest to the highest resolution HierCC level, a predicted resistant isolate is assigned to only one resistant cluster at any given HierCC level. This approach ensures that resistant clusters contain non-overlapping sets of resistant isolates.

### Identification of TPC of resistant HierCC clusters

The temporal patterns of resistant HierCC clusters were determined using similar methods as described by Kaur *et al*. [53]. Briefly, the sampling frequency of each resistant cluster from a given antibiotic were normalised based on the average yearly count for that cluster. The normalization was performed in python (version 3.9.6). The resistant clusters to 15 antibiotics or antibiotic combinations were then clustered in TPCs based on similarities in temporal abundance patterns to determine the timing of their emergence and changes in sampling frequency over time. The optimal number of TPCs for each dataset was determined through a combination of the silhouette index and manual inspection of the temporal delineation of the clusters. The silhouette index was calculated using the R package Bios2mdS (version 1.2.3) [56]. TPCs were then generated using the unsupervised c-means clustering algorithm in the Mfuzz (version 2.60.0) package of R (version 4.3.1) [57].

### Hierarchical representations of resistant HierCC clusters

The hierarchical representation of resistant HierCC clusters were generated using python (version 3.9.6) scripts from AMR-analysis [53]. Visualisation of the hierarchical representations were performed using graphviz’s (version 8.0.5) hierarchical layout engine [58]. Human and animal host information from the NCBI database was manually simplified to represent the closest associated animal, and the top five hosts of the isolate within each cluster were displayed.

## Results

### *C. jejuni* global dataset

We curated a *C. jejuni* global dataset of 63,102 genomes, of which 78.68% (n = 49,653) had country information, 67.08% (n = 42,347) year of collection, and 64.96% (n = 40,996) both year and country of collection. Genomes with both the year and country information were collected from 44 countries between 1905 to 2023, with 49.89% (n = 31,483) of the genomes coming from the USA and UK. However, surveillance data from UK ceased from 2020 onwards in the NCBI SRA database for unclear reasons, therefore, analysis of the UK dataset focused on between 2016-2019 data (**Supplementary Figure 1**). The UK and USA initiated WGS surveillance for *C. jejuni* from 2016-19 and 2018-2022, respectively, and the USA had collected substantial amount of *C. jejuni* WGS genomes (n=6,051) prior to the initiation from 2016 to 2017. Three datasets were constructed for downstream analysis. The UK dataset included genomes from two timeframes: 2016-17 and 2018-2019. The USA dataset consisted of genomes from three timeframes: 2016-17, 2018-19, and 2020-2022. Lastly, the “Other” dataset with genomes collected from countries outside the UK and the USA from three timeframes: 2016-17, 2018-19, and 2020-2022.

### Development and characteristics of *C. jejuni* multilevel HierCC typing scheme

The UNSW *C. jejuni* cgMLST scheme core genome consisted of 1161 core loci and a combined nucleotide length of 1.07 Mbp, which is 66.8% of the *C. jejuni* reference NCTC11168 genome length. Adjusted rand index (ARI) was calculated between MLST STs and clusters at each HC level to determine which HierCC level (HC level) provides clustering most similar to the pubMLST MLST scheme [50] (**Supplementary Figure 2**). HC264 had the highest ARI with the pubMLST MLST scheme and was set as the lowest resolution level for the multilevel HierCC typing scheme. The ARI between two nearby HierCC levels from HC0 to HC264 were calculated to identify “stable blocks” of HierCC clustering (**Supplementary Figure 3**). Similar to HCCeval, “stable blocks” were defined as sets of HierCC levels that define highly similar clusters (ARI >= 0.97) [39]. Within each stable block, the HierCC level with the highest Silhouette score with the cgMLST allelic profile distance matrix was identified. The final multilevel HierCC typing scheme comprises nine levels with increasing typing resolution: HC264, HC167, HC100, HC55, HC19, HC12, HC6, HC4, and HC2 (**Table 1**).

**Table 1.** Characteristics of the *C. jejuni* multilevel HierCC typing scheme.

| HierCC level | Number of clusters | Percentage increase of clusters from the lower resolution | Size of largest cluster (no. of genomes) | Average size of cluster | Number of singleton clusters |
| --- | --- | --- | --- | --- | --- |
| HC2 | 44148 | 20% | 338 | 1 | 38152 |
| HC4 | 36785 | 17% | 401 | 2 | 30228 |
| HC6 | 31500 | 38% | 521 | 2 | 24949 |
| HC12 | 22794 | 29% | 877 | 3 | 16975 |
| HC19 | 17726 | 112% | 1142 | 4 | 12729 |
| HC55 | 8362 | 66% | 1355 | 8 | 5369 |
| HC100 | 5036 | 63% | 2250 | 13 | 2955 |
| HC167 | 3099 | 60% | 3203 | 20 | 1640 |
| HC264 | 1935 | N/A | 4752 | 33 | 982 |

### Application of multilevel HierCC typing to track international movements of *C. jejuni*

The multilevel HierCC typing scheme revealed a highly diverse population structure in the *C. jejuni* global dataset (**Table 1**). Notably, the largest cluster at HC264 (cluster 105) accounted for only 7.53% of all isolates. Among the major clusters (clusters with more than 10 genomes assigned), continent-specific clusters were identified with more than or equal to 85% of the genomes from one continent. HC19 was selected as the highest resolution level to examine country- and continent-specific clusters as it is the broadest HierCC level where more than 90% of genomes were within country-specific clusters (**Supplementary Table 2**). Different clusters exhibited distinct global distribution patterns and more detailed country associations within the broader cluster can be revealed by further subdividing genomes at higher-resolution HierCC levels. We used three clusters in HC264 (cluster 27, 275, and 115) to demonstrate the utility of the scheme to track international movements of *C. jejuni* strains (**Figure 1**). For example, HC264 cluster 115 has a North America specific distribution, consisting predominantly of isolates from the USA and only 2 isolates collected from Canada. Internationally distributed HC264 cluster 275 can be subdivided into cluster 1142 and cluster 2961 at HC100, which were predominantly from the USA and Italy respectively. HC264 cluster 27 is a globally distributed cluster comprising 2337 genomes. Within this cluster, isolates can be sub-divided into two continent-specific clusters at HC167: HC167 cluster 27 for Europe and HC167 cluster 103 for North America. At HierCC 100, the continent specific clusters could be dissected further and identify country-specific clusters for the USA (HC100 cluster 103 and HC100 cluster 125), and the UK (HC100 cluster 27). A globally distributed HC100 cluster 11975 was also identified with 83 genomes. Most of the genomes were recovered from European countries (Germany, France, the Netherlands, Luxembourg, Denmark and the UK), and a smaller number of genomes from the USA (n=3), Egypt (n=2), and Chile (n=1).

**Figure 1.**
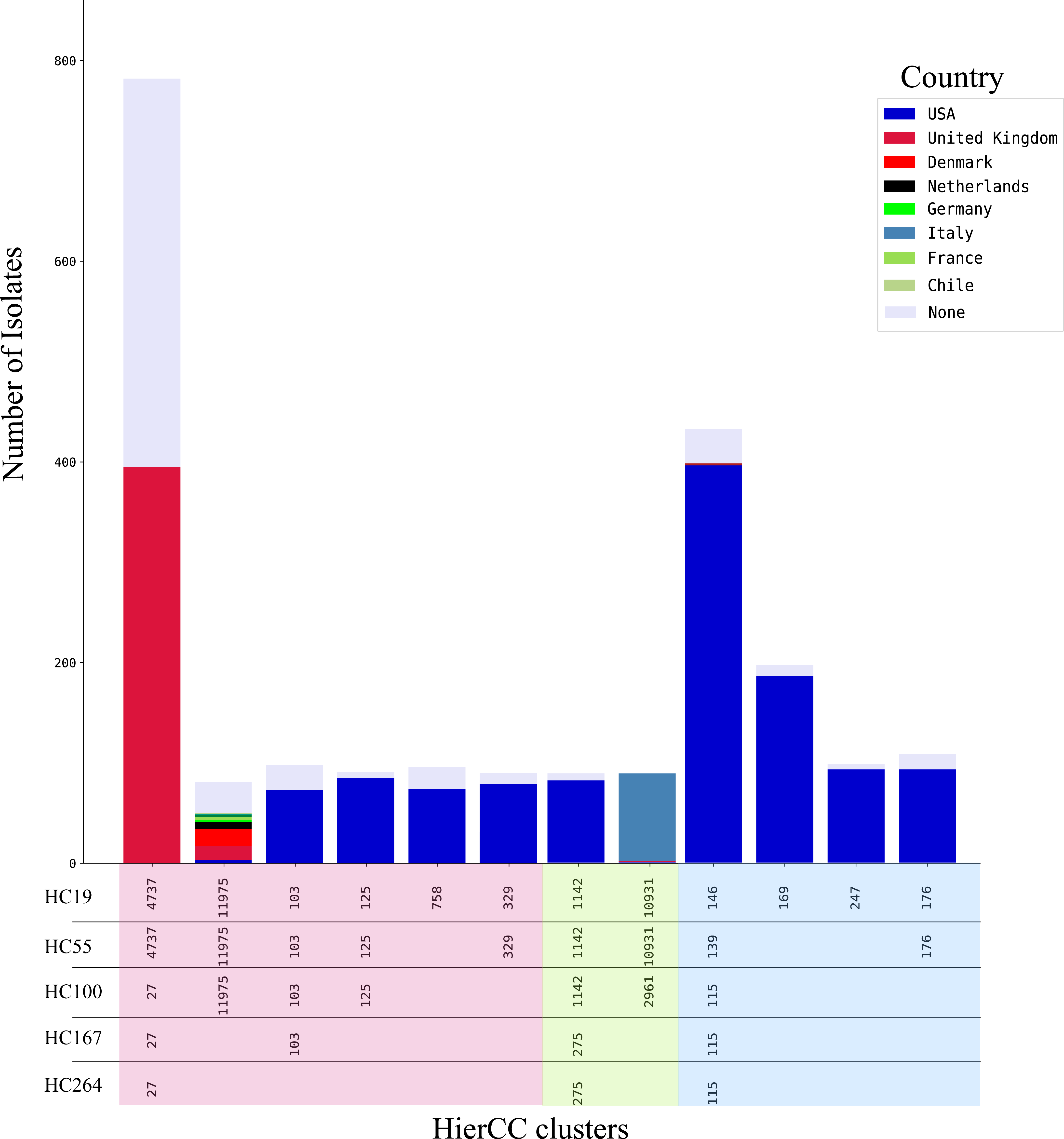
Geological diversity of *C. jejuni* in HC264 clusters 27, 275, and 115. The isolates in each column are denoted with country of origin as shown in the legend. Each column represents a unique HC19 cluster. The columns are arranged to reflect relatedness across HC264 to HC19, with columns most related to each other at the lower resolution levels appearing closer. The HierCC clusters are coloured based on which HC264 cluster they belonged to in the following order: HC264 cluster 27 is coloured in light red, HC264 cluster 275 is coloured in light green, and HC264 115 is coloured in light blue. For example, from left to right, column 1-6 all belonged to cluster 27 at HC264. At HC167, column 1 and 2 both belonged to HC167 cluster 27, and column 3 to 6 all belonged to HC167 cluster 103.

### Application of multilevel HierCC typing to host attribution and transmission

Metadata analysis was performed to identify potential association between the host types and HierCC clusters. In total, 41,076 (65.1% of the total global dataset) genomes had host information and were from 49 different hosts. Among these genomes, human samples constituted 43.93% (18,047 genomes) of the genomes whereas 53.7% (22057 genomes) were collected from just five farm animal hosts: chicken (35%, 14,718 genomes), cattle (15.31%, 6,289 genomes), sheep (1.18%, 486 genomes), turkey (0.71%, 293 genomes), and swine (0.66%, 271 genomes). A farm-animal dataset was built consisted of all 22057 genomes from chicken, cattle, sheep, swine, and turkey. In this dataset, farm-animal-associated clusters are defined as clusters with more than 90% of the isolates from one type of farm animal. Using this definition, we identified HC55 as the highest resolution level to examine farm-animal-associated clusters, because majority (≥ 70%) of all genomes are within farm-animal-associated clusters at this level (**Supplementary Table 3**). The farm-animal dataset was combined with 18,047 human clinical genomes to identify potential host associations to human infections.

In this human and farm animal dataset (n=40,104), majority of the HC264 clusters consisted of isolates from multiple host species indicating a generalist lifestyle among lineages that infect human [59]. We selected HC264 cluster 30 and 32 to demonstrate the utility of the multilevel HierCC typing for farm-animal source attribution (**Figure 2**). HC264 cluster 30 showed a host preference between human and predominately cattle. Further examination at HC100 revealed the existence of two human-cattle-sheep clusters (cluster 30, and 191) and two human-cattle clusters (cluster 183 and 190) and one human-cattle-swine clusters (cluster 144).

**Figure 2.**
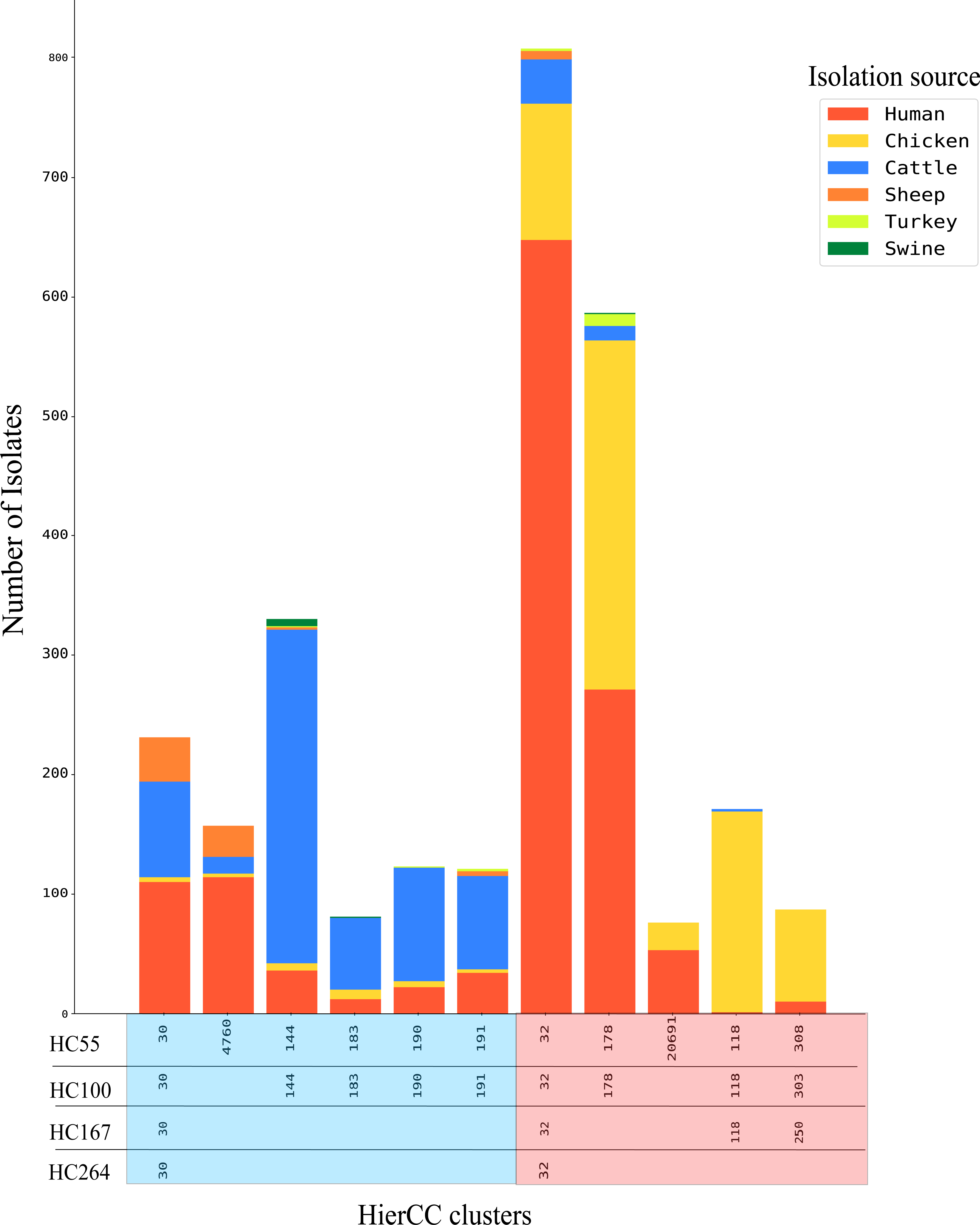
Host diversity in HC264 cluster 30 and 32. The isolates in each column are denoted with isolation source as shown in the legend. Each column represents a unique HC55 cluster. The columns are arranged to reflect relatedness across HC264 to HC55, with columns most related to each other at the lower resolution levels appearing closer. The HierCC clusters are coloured based on which HC264 cluster they belonged to in the following order: HC264 cluster 30 is coloured in light blue, HC264 cluster 32 is coloured in light red. For example, from left to right, column 1-6 all belonged to cluster 30 at HC264 and At HC167, column 1 and 2 both belonged to H100 cluster 30, and column 3 to 6 all belonged to a unique HC100 cluster.

HC264 cluster 32 is a generalist cluster with 2101 genomes, of which 55.26% were from human and 39.08% were from chickens. By further partitioning the genomes at HC167, we identified HC167 cluster 32 and 250 with hosts of both human and chicken, and cluster 118 with chicken as the only host. Of the 1613 HC167 cluster 32 genomes with host information, 65.34% (1054 genomes) were from humans and 28.02% (452 genomes) from chickens.

HC167 cluster 250 had 75.5% (145 genomes) from chickens and 20.85% (39 genomes) from humans. This contrasted with the chicken specialist HC167 cluster 118, where 98.29% (172 genomes) of the genomes were from chickens and only one (0.5%) from humans. At HC55, HC167 cluster 32 can be further subdivided into three clusters (HC55 cluster 32, 178, and 20691). HC55 cluster 32 and 178 infect predominantly humans and chickens but also other farm animals and HC55 cluster 20691 infects only humans and chickens. HC55 cluster 32 and 178 are generalist clusters infecting multiple animal hosts sharing the farm environment and humans. HC55 cluster 20691 is similar to HC55 cluster 308 in that they were only found in chickens and humans.

### Resistance score of global *C. jejuni* isolates

Using a dataset of 3,181 genomes with phenotypic antibiotic susceptibility information from the NCBI pathogen detection database (https://www.ncbi.nlm.nih.gov/pathogens), we assessed the accuracy of AbritAMR predicting phenotypic resistance of 8 antibiotics (ampicillin, ciprofloxacin, tetracycline, azithromycin, erythromycin, gentamicin, nalidixic acid, and telithromycin) (**Supplementary Table 4**). Overall, AbritAMR had an accuracy of 97.77% for interpreting phenotype resistance from genomic determinants in *C. jejuni*. Based on the prediction of AbritAMR to a total of 15 antibiotic classes or combinations, we assigned resistance score from 0 to 15 to each genome in the global dataset. The most common resistance determinants of each geographic location for narrow-spectrum beta-lactams, tetracyclines, quinolones and macrolides for different countries and timeframes are shown in **Supplementary Table 5**. The complete list of resistance determinants detected by AbritAMR for the 15 antibiotic classes or combinations are shown in **Supplementary Table 6**.

A detailed overview of resistance scores in the global dataset across the USA, the UK, and other countries between 2016 and 2022 is presented in **Supplementary Table 7**. In total, only 8.27% of the isolates were predicted to be susceptible to all the antibiotics and majority (46.58%) of the isolates were predicted to be resistant to only one drug class. In the USA, the overall resistance remained stable and decreased 1.91% from 2016-2017 to 2018-2019 and decreased further by 1.81% from 2018-2019 to 2020-2022. Resistance to one antibiotic class has increased significantly by 4.7% in the USA from 2016 to 2022. In the UK, the overall resistance percentage in the UK remained stable from 2016-2017 to 2018-2019, but the average proportion of susceptible isolates was lower (∼3.5%) compared to the USA (∼12%) in the same time period. The resistance score also allowed identification of isolates resistant to multiple antibiotics and antibiotic classes in the global dataset (**Supplementary Table 7**). In total, 24.4% of the isolates were resistant to two drug classes, 17.98% of the isolates were resistant to three drug classes and 1.83 % of the isolates resistant to four drug classes.

### Resistance to medically important drug classes

In the global dataset, *C. jejuni* resistance to individual antibiotics varied (**Supplementary Figure 4**). Most notably, in descending order, 80.3% were resistant to narrow-spectrum beta-lactams, 40.4% to tetracyclines, 29.8% to quinolones, 3.4% to streptomycin and 1.2% to macrolides. From 2016-2017 period to 2020-2022 period, the global resistance to narrow-spectrum beta-lactams, tetracyclines and quinolones have reduced significantly (*p* < 0.0008) (**Figure 3**). Resistance to amikacin, kanamycin, and streptomycin have increased significantly from 2018-2019 period to 2020-2022 period.

**Figure 3.**
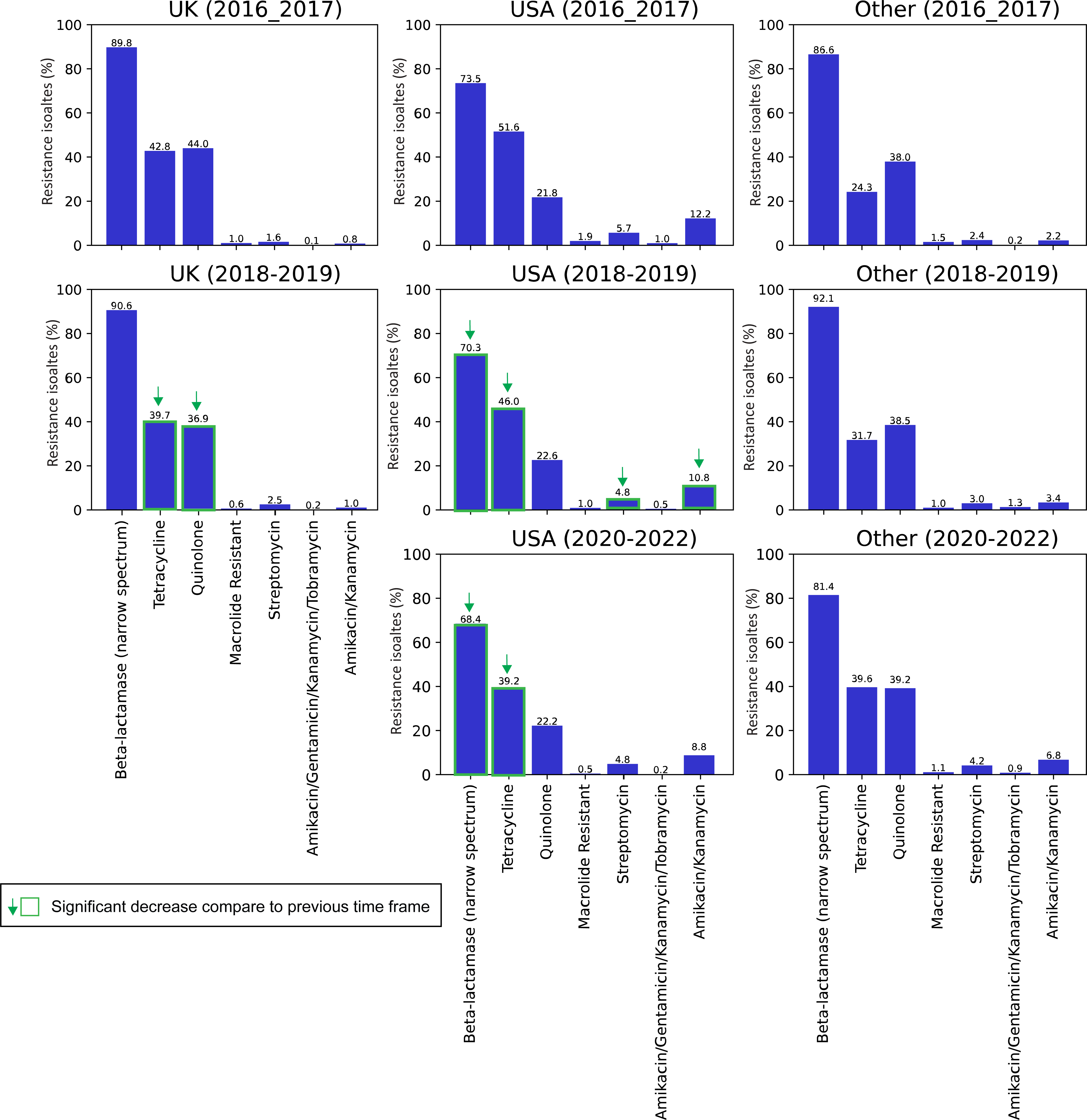
Resistant isolates to seven antibiotic classes or antibiotic combinations the UK, the US and other countries. Percentage of resistant isolates to seven antibiotic classes or combinations within the local datasets of available times frames. Resistance to a given antibiotic category in different time frames was compared using the binomial test. Significance was assessed at the Bonferroni corrected *p* < 0.0008. Significant decrease in resistance is indicated by a downwards green arrow.

The resistance rate to medically important antibiotics in the UK, the USA and other countries are shown in **Figure 3**. In the UK from the 2016-2017 period to the 2018-2019 period, resistant to tetracyclines and quinolones decreased significantly. Resistance to narrow-spectrum beta-lactams remained high with no significant change during the same time frame. In the USA, resistant to narrow-spectrum beta-lactams, tetracyclines, amikacin, kanamycin and streptomycin all significantly decreased from the 2016-2017 period to the 2018-2019 period. The decreasing trend for narrow-spectrum beta-lactams and tetracycline continued from the 2018-2019 period to the 2020-2022 period.

### 50s_L22 mutations in the global dataset

Although the percentage of isolates with predicted phenotypic resistance to macrolides is low in the global dataset, we identified significant increases of macrolides induced mutations in the *rplV* gene, coding for the 50S ribosomal protein L22: *50S_L22 (A103V)* and *50S_L22 (G86E)*. In the global dataset, there was a 15% increase of genomes carrying *50S_L22* gene mutations from 2016 to 2022 (**Figure 4**). In the USA dataset from 2016-2022, 27% (n =5866) of genomes carried *50S_L22* mutations and there was an 15.5% increase of genomes carrying *50S_L22* mutations from the 2016-2017 period to the 2020-2022 period. We also examined the relationship between phenotypic resistance to macrolides (azithromycin and erythromycin) and the presence of both *50S_L22 (A103V)* mutation and *CmeABC* gene. We found 560 isolates with both *50S_L22 (A103V)* mutation and *CmeABC* gene and phenotype resistance data in the global dataset, all of which are susceptible to both azithromycin and erythromycin.

**Figure 4.**
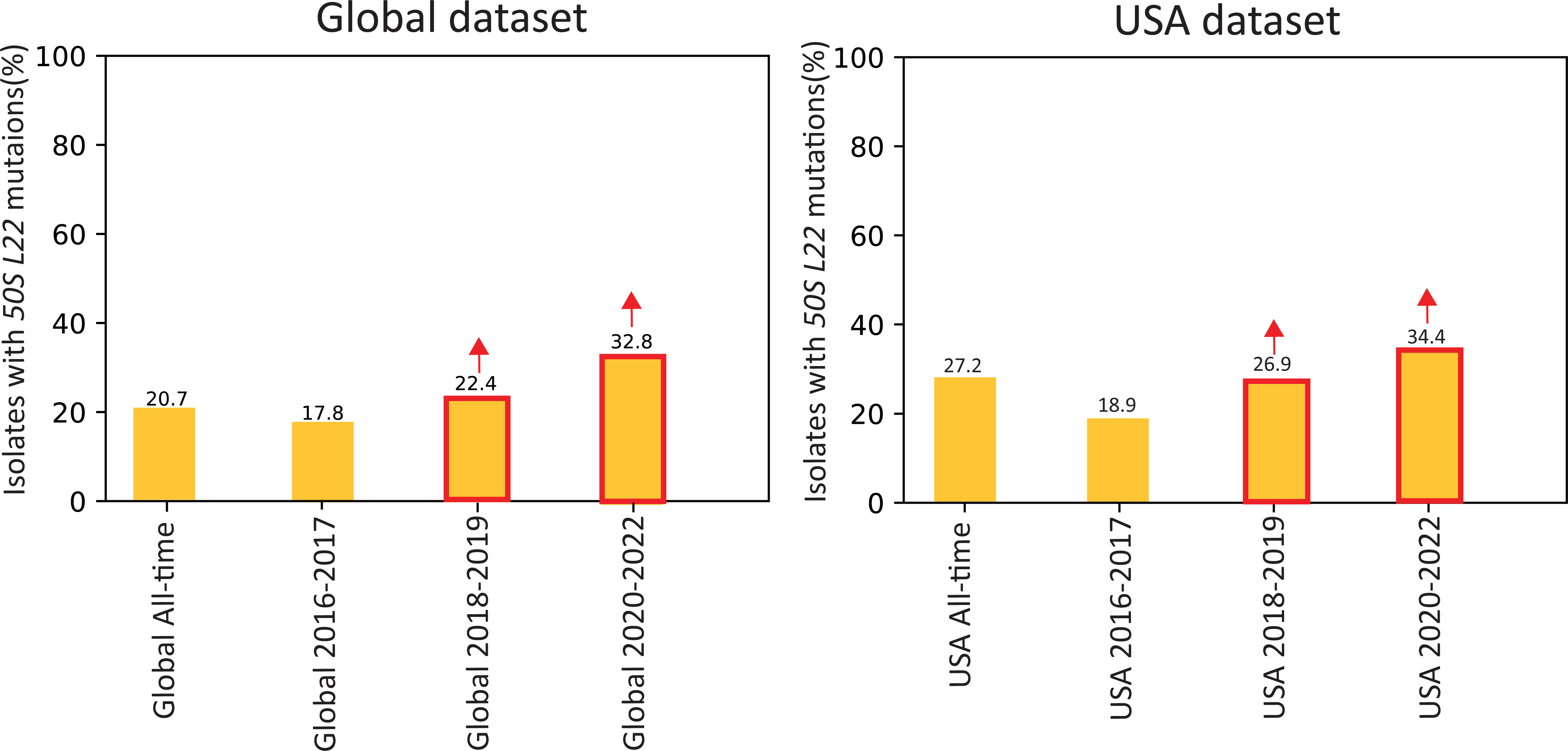
***50S_L22* mutations in the global and USA dataset**. Percentage of isolates with *50S_L22* mutations within the datasets of available times frames. Isolates carrying the mutation in different time frames was compared using the binomial test. Significance was assessed at the Bonferroni corrected *p* < 0.0008. Significant increase in between time frames is indicated by an upwards red arrow. Percentage of isolates in both the global and USA dataset has increased significantly from 2016-17 period to 2018-2019 period, and again in 2020-2022 period

### Application of multilevel HierCC typing to identify resistant HierCC clusters and AMR surveillance

A “resistant HierCC cluster” for a given antibiotic was defined as a HierCC cluster in which at least 80% of the isolates were predicted to be resistant to that antibiotic. To ensure mutual exclusivity between resistant clusters between different HierCC levels, each resistant isolate was accounted for only one cluster at any HierCC level. Large clusters with distinctive resistance patterns can then be further dissected at higher resolution HierCC levels to explore intricate links between AMR and HierCC clusters. The USA dataset was chosen for the resistant HierCC cluster analysis as it is the largest dataset with genomes collected from 2016 to 2022.

### Resistance HierCC clusters to medically important antibiotics in the USA from 2016-2022

The USA dataset from 2016 to 2022 was chosen to showcase the potential utility of the multilevel HierCC scheme for nation-wide AMR surveillance. A total of 276 resistant clusters were identified in the USA from 2016-2022, of which 206 resistant HierCC clusters were identified for narrow-spectrum beta-lactams, tetracyclines, quinolones and macrolides (**Supplementary table 8**). We normalised the yearly abundance of resistant clusters against the average yearly count for each resistant cluster. This allowed us to group resistance clusters with similar temporal trends into temporal pattern clusters (TPCs), regardless of the size of the resistance cluster. We identified historical and emerging resistant clusters from the USA dataset between 2016 to 2022. Nine distinctive TPCs were identified from resistant clusters of all antibiotic classes in the USA dataset from 2016 to 2022 (**Figure 5**). The TPCs have been arranged in chronological orders of when the peak abundance occurred. There were 24 resistant clusters in TPC 1, all of which contained 2-fold more isolates in 2016 compared to other years and the frequency had reduced steadily in more recent years. TPC 2 to 7 showed groups of resistant clusters with abundance peaked at various points between 2017-2020, but frequency reduced significantly in more recent years. Resistant clusters in TPC 8 and 9 showed a steep rise in abundance between 2021 and 2022, indicating emerging resistant clusters potentially associated with recent outbreaks. The temporal trends shown in **Figure 5** examined the overall resistance trend of all 15 antibiotics and classes.

**Figure 5.**
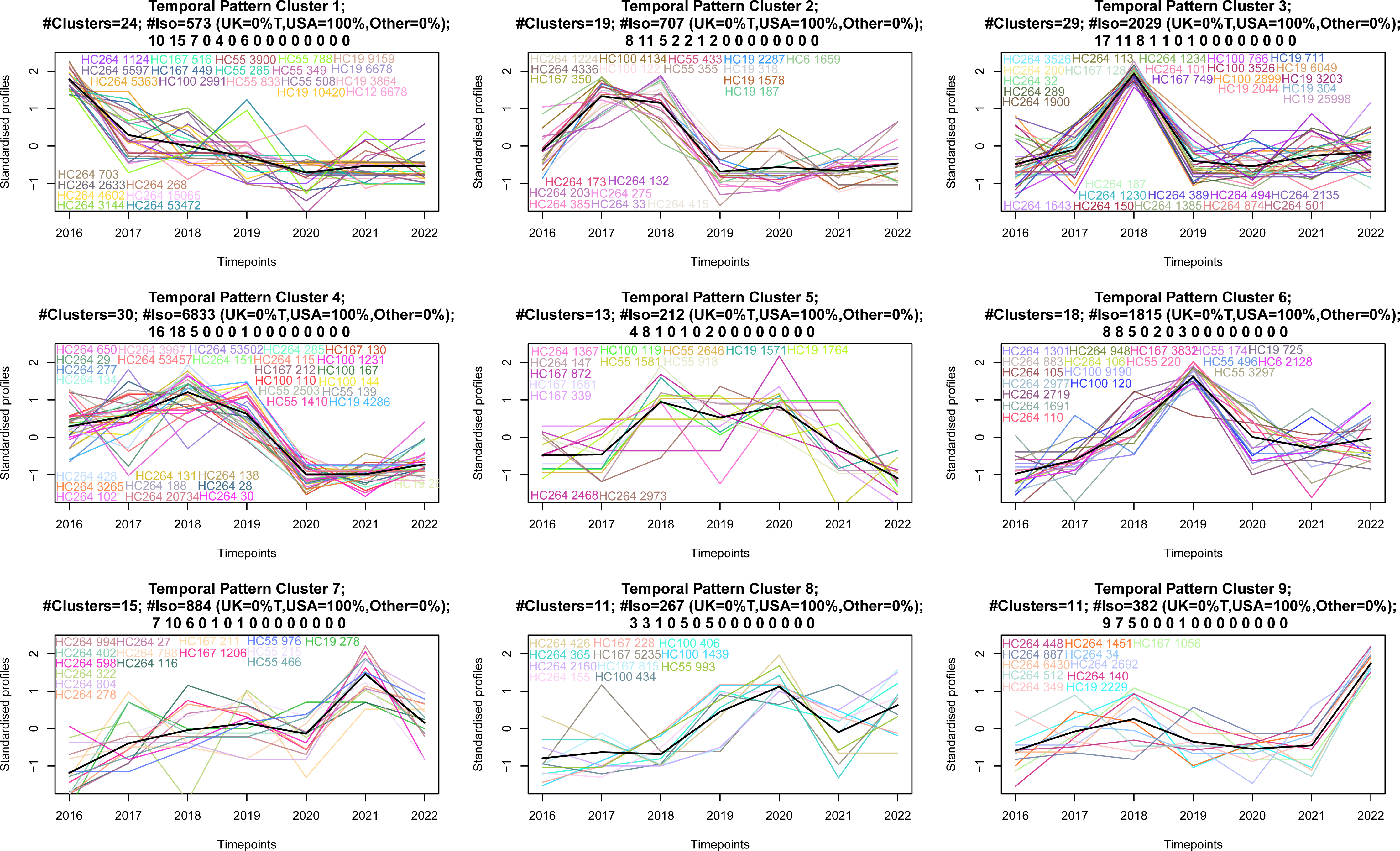
Temporal resistant pattern of 15 antibiotic classes or combinations in the USA in 2016-2022. Temporal trends of resistant HierCC clusters between 2016-2022 in the USA dataset. Nine TPC contained 170 resistant clusters to 15 antibiotic classes or combinations. An individual graph with separate header was created for each TPC. Each header indicates the number of resistant clusters (#clusters) within the TPC, total number of isolates (#Iso) and the country of origin of the isolates (UK, USA or other). The series of numbers at the bottom of the header indicated the number of HierCC clusters resistant to a particular antibiotic class or combination. (e.g.: 10 15 7 0 4 0 6 0 0 0 0 0 0 0 0 in the first graph, here 10 means 10 of the 24 clusters in this TPC are resistant to narrow-spectrum beta-lactamase). The order of the 15 antibiotic classes or combinations are as follows: 1. Beta-lactamase (narrow-spectrum), 2. Tetracycline, 3. Quinolone, 4. Macrolide, 5. Streptomycin, 6. Amikacin/Gentamicin/Kanamycin/Tobramycin, 7. Amikacin/Kanamycin, 8. Kanamycin/Tobramycin, 9. Amikacin/Kanamycin/Tobramycin, 10. Lincosamides, 11. Lincosamide/Streptogramin, 12. Chloramphenicol, 13. Chloramphenicol/Florfenicol, 14. Sulfonamide, 15. Florfenicol/Oxazolidinone.

### Narrow-spectrum beta-lactams

Temporal trends of beta-lactam resistant clusters in the USA from 2016-2022 are shown in **Supplementary Figure 5**. Overall resistance to beta-lactams in the USA decreased by 5.1% from the 2016-17 period to the 2020-2022 period (**Figure 3**). The decrease in overall resistant correlated to the temporal trends in TPC 1-7, where sampling frequency of 67 resistant clusters in these TPCs fell below average in the 2020-2022 period. Despite overall decrease in the percentage of beta-lactam resistant strains in the USA dataset from the 2016-17 period to the 2020-2022 period, 15 resistant clusters in TPC 8-9 showed on average 1.5 to 2-fold increase in frequency in the 2020-2022 period.

### Tetracyclines

Temporal trends of tetracycline resistant clusters in the USA from 2016-2022 are shown in **Supplementary Figure 6**. TPC 1-7 representing 69 tetracycline resistant clusters had reduced frequency in the 2020-2022 period, falling below average. Resistant clusters in TPC 8 showed small increase in frequency since 2020 and peaked at 1-fold increase in 2021. TPC 9 had 7 resistant clusters with 2-fold increase in frequency in 2022.

### Quinolones

Temporal trends of quinolone resistant clusters in the USA from 2016-2022 are shown in **Supplementary Figure 7**. Percentage of isolates resistant to quinolones remained stable in the USA from 2016-2022 (**Figure 3**). Sampling frequency of TPC 3, 4 and 5 peaked at year 2018, 2019 and 2021 respectively, and TPC 6 with 5 resistant clusters showed 2-fold increase in sampling frequency in 2022.

### Hierarchical identification of resistant clusters in the USA from 2016-2022

The *C. jejuni* population in the US can be divided into many sporadic HierCC clusters at the broadest HierCC level HC264, which was expected as *C. jejuni* population is highly diverse [60]. Hierarchical networks of resistant clusters were created to identify acquisition of resistance in the USA dataset, which provided novel insights into *C. jejuni* antimicrobial resistance acquisition. Four cases of hierarchical networks are presented below (**Figure 6)**. Scenario A showcased emergence of resistant clusters to one drug at higher resolution HierCC levels inside a non-resistant cluster at a lower resolution level. HC264 cluster 112 is not a resistant cluster to tetracyclines, but at higher resolution levels (HC167 and HC100), it contained two tetracyclines resistant clusters, which are HC167 cluster 815 and HC100 cluster 120 (**Figure 6A**). HC167 cluster 815 was mostly isolated from chickens and humans and HC100 cluster 120 was mostly isolated from chickens and dogs. Scenario B showcased emergence of resistant clusters to multiple antibiotics at higher resolution HierCC levels inside a non-resistant cluster at low resolution level. HC264 cluster 111 is a non-resistant cluster found only in chickens but contained three resistant clusters to multiple antibiotics at HC167 and HC100 (**Figure 6B**). HC167 cluster 872 was only resistant to beta-lactams. HC167 cluster 1206 was resistant to tetracyclines, streptomycin, amikacin and kanamycin. HC100 cluster 434 was resistant to streptomycin, amikacin and kanamycin. In Scenarios C and D, clusters resistant to one drug class at lower resolution HierCC levels were found to have acquired additional resistance to more antibiotics at higher resolution levels. HC264 cluster 105 was resistant to beta-lactams and found in human, chicken, cattle, turkey and wild birds (**Figure 6C**). Within HC264 cluster 105, HC100 cluster 406 acquired multi-drug resistance to beta-lactams, tetracyclines, and quinolones and was found only in chickens; and HC55 cluster 976 acquired additional resistance to beta-lactams and tetracyclines. Similarly, HC264 cluster 115 was resistant to beta-lactams only and found in cattle, chickens, humans, turkey, and sheep (**Figure 6D**). It contained two multi-drug-resistant clusters to beta-lactams, tetracyclines and quinolones at HC55 and HC19. The two higher resolution clusters were found in different hosts, with HC19 cluster 711 found in cattle and chickens, and HC55 cluster 139 was found in cattle, cattle, chickens, human, turkey, and sheep.

**Figure 6.**
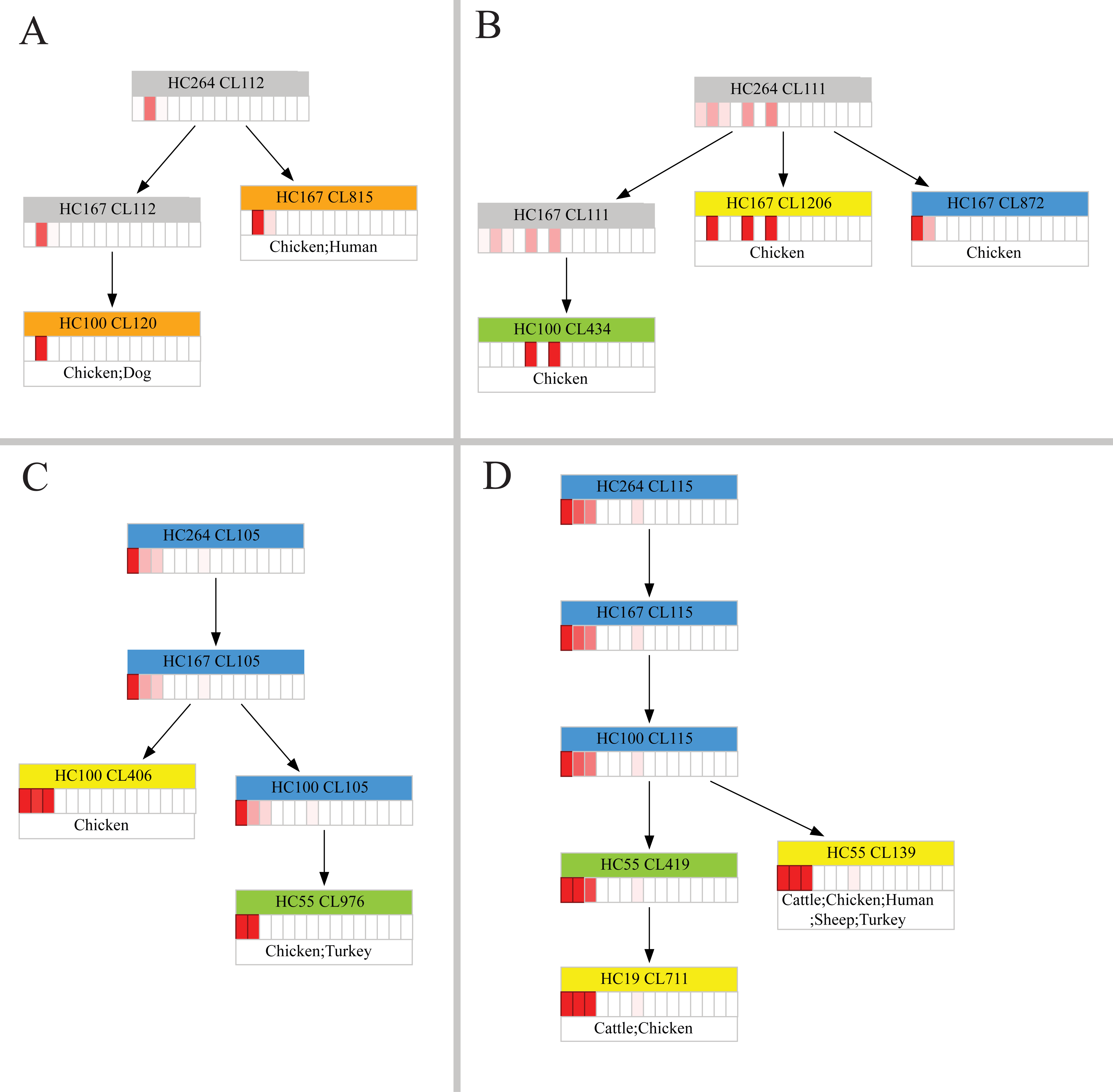
Hierarchical identification of resistant HierCC clusters in the USA from 2016-2022. Four scenarios of hierarchical identification of resistant HierCC clusters with at least 10 isolates in the USA from 2016 to 2022. None-resistant cluster were coloured in grey and resistant clusters are coloured based on the corresponding drug in the following order: narrow-spectrum beta-lactams (blue) and tetracyclines (orange). Clusters resistant to two antibiotics are coloured in green and clusters resistant to three antibiotics are coloured in yellow. **A**. Shows two tetracycline resistant clusters. **B**. Shows two streptomycin resistant clusters and one narrow-spectrum beta-lactams resistant clusters. **C**. Shows two tetracycline resistant clusters and a series of narrow-spectrum beta-lactams resistant clusters. **D**. Shows two quinolones resistant clusters and a series of narrow-spectrum beta-lactams resistant clusters. The heatmap is placed at the bottom of each resistant cluster to indicate the percentage of resistant isolates to antibiotics in the following order: 1. beta-lactamase (narrow-spectrum), 2. tetracycline, 3. quinolone, 4. macrolide, 5. streptomycin, 6. amikacin/gentamicin/kanamycin/tobramycin, 7. amikacin/kanamycin, 8 kanamycin/tobramycin, 9. amikacin/kanamycin/tobramycin, 10. lincosamides, 11. lincosamide/streptogramin, 12. chloramphenicol, 13. chloramphenicol/florfenicol, 14. sulfonamide, 15. florfenicol/oxazolidinone. A black outline indicated at least 80% of the isolates were non-susceptible to the antibiotic at that rectangle. Beneath the heatmap, the most frequent isolation sources for the isolates in the cluster are listed.

### Temporal trends of *50s_L22* mutation in response to macrolides

Considering that macrolides (azithromycin and erythromycin) are medically important antibiotics, tracking of temporal trends in *50s_L22* mutations is crucial for identifying and monitoring underlying selection pressure. To achieve this, “*50S_L22* mutation HierCC cluster” was defined as a HierCC cluster in which at least 80% of the genomes were carrying *50S_L22* mutations. Temporal patterns of *50S_L22* mutation clusters in the USA from 2016-2022 are shown in **Figure 7**. Frequency of *50S_L22* mutation clusters in TPC 1-3 peaked prior to 2020 and gradually decreased afterwards. Frequency of *50S_L22* mutation clusters in TPC 4-5 has peaked prior to the 2020-2022 period and remain stable in more recent years. TPC 6 contained three *50S_L22* mutation clusters with 1.5-fold increase in frequency between 2020-2022.

**Figure 7.**
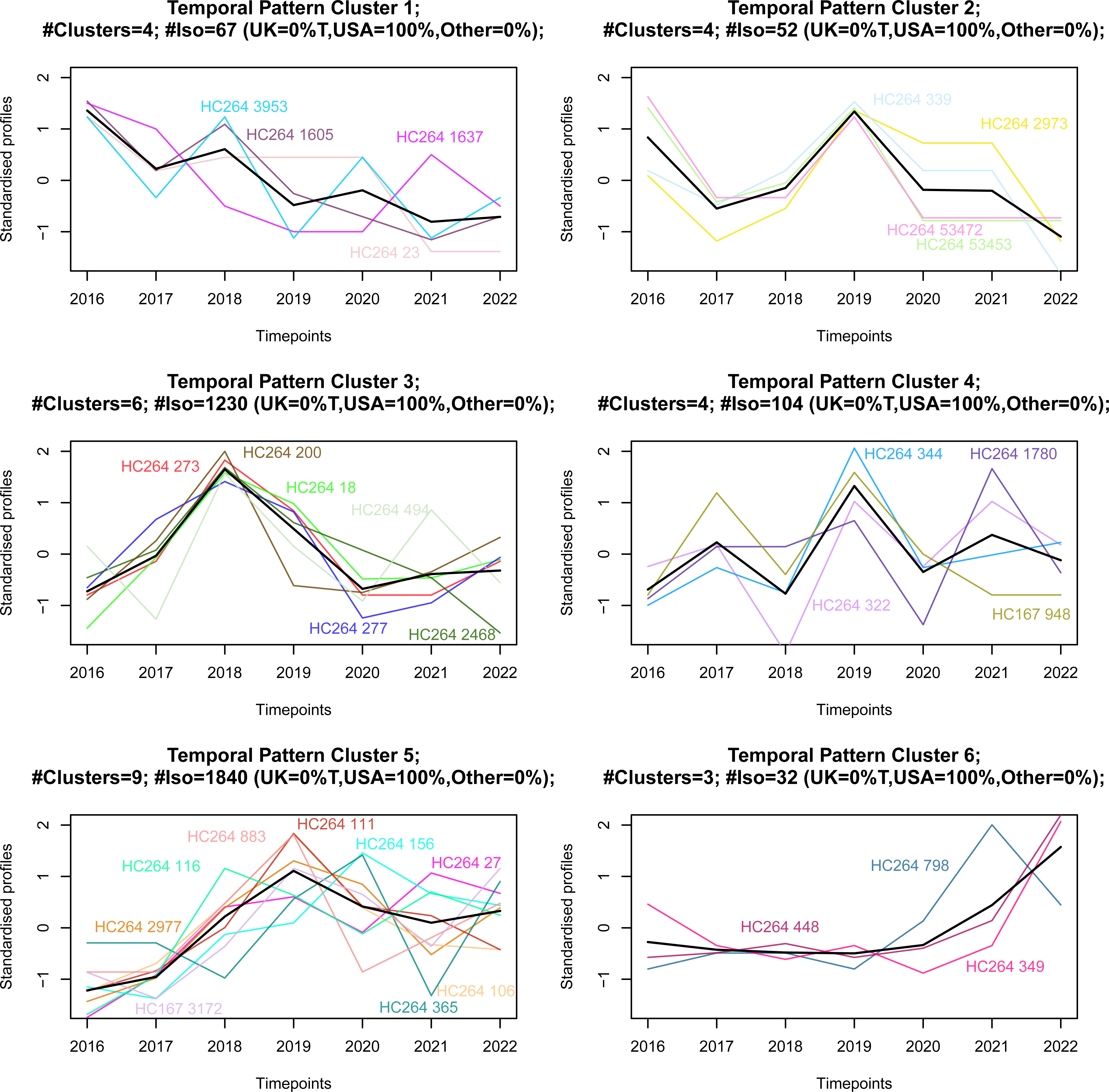
Temporal pattern of *50S_L22* mutations in the USA in 2016-2022. Temporal trends of *50S_L22* mutation HierCC clusters between 2016-2022 in the USA dataset. Six TPCs contained 30 *50S_L22* HierCC clusters. For each TPC, temporal profile of each unique HierCC cluster is the standardised count of isolates per year, which was represented by a unique coloured line. The weighted average temporal pattern of all clusters in a TPC is represented by a bold-black line. An individual graph with separate header was created for each TPC. Each header indicates the number of resistant clusters (#clusters) within the TPC, total number of isolates (#Iso) and the country of origin of the isolates (UK, USA or other).

## Discussion

A stable and standardised genotyping scheme is essential for accurate epidemiological monitoring and AMR surveillance of *C. jejuni*. In this study, we developed a novel multilevel HierCC typing scheme to provide standardised classification for publicly available *C. jejuni* WGS genomes worldwide (n= 63102 as of June 2023). The multilevel design provided flexible typing resolutions comparable to traditional MLST and cgMLST and enables the selection of appropriate typing resolutions for both short- and long-term epidemiological investigations. We used publicly available surveillance genomic data to demonstrate the utility of the *C. jejuni* multilevel HierCC typing scheme for tracing global transmission, identifying host sources, and monitoring AMR trends.

### Comparison to the Oxford cgMLST scheme

Cody *et al.* designed an open access PubMLST *C. jejuni*/*coli* cgMLST scheme consisting of 1343 loci in v1.0 and 1142 loci in v2.0 [50]. The system was defined using both *C. coli* and *C. jejuni* strains from Oxfordshire, UK between 2011 and 2014 [50]. The UNSW *C. jejuni* cgMLST scheme was developed using *C. jejuni* isolates representing 39 countries and 2587 MLST STs. The resulting core gnome consisting of 1161 loci, which is smaller than v1.0 and larger than the v2.0 of the Oxford cgMLST scheme. The core genome of the UNSW scheme as expected, was larger than v2.0 of the Oxford scheme, as the representative dataset was comprised exclusively of *C. jejuni* genomes. The UNSW *C. jejuni* cgMLST scheme provides a species-specific typing scheme that captures the global diversity of *C. jejuni*.

### Epidemiologically meaningful HierCC levels selected for epidemiological applications

The European Society of Clinical Microbiology and Infectious Disease published a guideline for developing novel classification technologies for bacterial epidemiology in 2007 [61]. These guidelines underline the importance of developing new typing methods that leverage readily available whole gnome data more cost-effectively. Prior to this study, hierarchical clustering of cgMLST with arbitrarily determined allelic difference levels had been applied to small *C. jejuni* datasets for AMR detection (n=511) or animal source attribution (n=622) [62,63]. Most notably, Hsu *et al*. applied the Oxford cgMLST scheme to 622 *C. jejuni* isolates and used six levels of allelic difference of 5, 10, 25, 50, 100, and 200 and demonstrated its utility to identify food animal source for human infections [63]. In this study, we curated a large global dataset (n=63102) of *C. jejuni* WGS genomes and identified nine epidemiologically meaningful HierCC levels (HC2, HC4, HC6, HC12, HC19, HC55, HC100, HC167, and HC264) using a statistical approach similar to HCCeval [39]. The epidemiologically meaningful levels provided a standardised recommendation of core genome allelic distances for surveillance applications, which would allow standardised communications and comparisons across time and regions. By integrating the nine levels into one comprehensive multilevel scheme, the multilevel HierCC typing scheme provides a standardised hierarchy for related strains with flexible typing resolution. Strains are grouped into larger clusters at broader HierCC levels and smaller clusters as resolution increases (**Supplementary table 1**). Because clusters at different HierCC levels contained a hierarchical set of genomes, the higher resolution clusters could be examined in the context of their lower resolution clusters to provide epidemiological inferences. Broader HierCC levels from HC264 to HC19 are useful for investigating longer-term and global epidemiology, while higher resolution levels from HC12 to HC2 can provide similar discriminatory power to cgMLST schemes for outbreak detection and other short-term applications. We demonstrated the potential utility of the scheme for epidemiological surveillance at local and global levels. We identified farm-animal associated clusters up to HC55, country and continent specific clusters up to HC19, and utilised the hierarchical network between the HierCC levels to track emergence and persistence of AMR genotypes.

### Multilevel HierCC typing scheme allows tracking of *C. jejuni* global distribution

Large numbers of livestock and animal products are traded each year across the globe in the age of globalisation, facilitating the spread of foodborne pathogens [64]. By identifying country and continent specific clusters at each HierCC levels, the multilevel HierCC scheme allows genomes with sporadic occurrence to be grouped into larger, more informative clusters. This utilisation can potentially provide useful information for tracking the movement of pathogenic *C. jejuni* strains beyond borders and identify international outbreak clusters. HC264 cluster 275 contained genomes from routine surveillance between 2013 to 2022 from the USA and genomes from a single camylobacteriosis outbreak in Italy, 2018 [65]. At HC100, the Italian outbreak cluster (HC100 cluster 2961) could be differentiated from the US-specific cluster (HC100 cluster 1142) (**Figure 1**). The connection between the US-specific cluster to the Italian outbreak would have been overlooked by traditional single-level typing scheme with fixed resolution.

### Multilevel HierCC typing scheme provides flexible typing resolution for *C. jejuni* farm animal host attribution

Based on preference to a single host or multiple hosts, *C. jejuni* lineages can be categorised as specialist or generalist respectively [66]. However, methods used to identify host-associated *C. jejuni* groups are inconsistent between studies using conventional MLST or its clonal-complex based typing system [66,67]. Comparing to these conventional methods, the flexibility of the multilevel HierCC scheme provided the appropriate tying resolution to identify the most likely farm animal source of human infections at a much higher precision **(Figure 2)**. The cluster nomenclature remained consistent at each HierCC level, while the flexible typing resolution allows for differentiation between specialist and generalist genotypes. This could potentially provide more accurate source attribution and aid more targeted approach for prevention and control.

### Antibiotic resistance in the global dataset

In the complete global dataset, over 90% of the isolates were predicted to be resistant to at least one antibiotic. The high overall resistance rate in *C. jejuni* global population can potentially be linked to the heavy reliance of antimicrobials to maintain health and productivity during intensive farm-animal production [18,68–70]. Variations in regulations and legislation across countries can result in differing antibiotic selection pressures.

Therefore, it is essential to analyse the global dataset by country of origin. The UK and the USA introduced WGS surveillance program for *C. jejuni* in 2016 and 2018, respectively [25,26]. These programs generated a large volume of high-quality WGS data in public domain, making them particularly valuable for AMR surveillance investigations.

In the UK, An AMR national action plan reduced the usage of antibiotics in humans and animals in the UK by 28% from 2014 to 2020 [71–73]. Correspondingly our analysis of the genome data found that while resistance to narrow-spectrum beta-lactams remained high, resistance to tetracyclines and quinolones significantly decreased from 2016–2017 to 2018– 2019.

In the USA, proportions of isolates resistant to beta-lactams and tetracycline has decreased significantly while resistance to quinolones remained stable in the USA from 2016-2022. Although resistance to macrolide remained low (∼2%) in both the UK and the USA, proportions of genomes with *50S_L22* mutations have increased significantly in the USA from 2016 to 2022. According to the FDA, the sale of medically important antibiotics for veterinary uses in the USA had increased by more than 8% from 2017 to 2020[19]. The widespread use of tylosin, a macrolide antibiotic, in cattle and chicken feed in the USA agricultural industry could be a factor contributing to the increase of *50S_L22* mutations [19,68–70,74,75].

For countries outside of the UK and the USA without a WGS surveillance program, the genomes were combined to show the prevalence of resistant strains in each time frames. Overall, predicted resistance in the other countries fluctuated from 2016 to 2022, which could be due to the sampling bias toward outbreak strains in these countries, as no countries other than the USA and the UK is actively monitoring *C. jejuni*. Although it is important to present the AMR prediction results for the other countries, we refrain from discussing the significance of these results due to the heterogenous nature of the dataset. This highlights the importance of consistent and continuous surveillance of *C. jejuni* and the need of a universal genotyping scheme that provides AMR prediction and tracking at global scale.

### Temporal trends and hierarchical network of resistant HierCC clusters to medically important antibiotics in the USA from 2016 to 2022

Using the USA 2016-2022 surveillance dataset, we demonstrated the utility of the multilevel HierCC typing scheme for identifying and monitoring of *C. jejuni* resistance to medically important antibiotic classes. *C. jejuni* can rapidly transfer resistance genes between strains via natural transformation or conjugation [76,77]. The multilevel HierCC typing scheme can provide the ideal typing resolution to identify clusters associated with antibiotic resistance. Each cluster can be consistently identified across time, allowing continuous monitoring of strains with specific resistance pattern. Strains with similar resistance patterns in broader levels often group together at higher resolution levels. The hierarchical structure enabled rapid identification of resistant clusters by sub-setting closely related strains with similar resistance pattern at higher resolution HierCC levels through network analysis.

We clustered the resistance clusters based on their temporal trend patterns. This application is particularly useful for pathogens with sporadic outbreak patterns such as *C. jejuni,* as not all resistance clusters are associated with recent outbreaks. In the USA data set, resistance to beta-lactams and tetracyclines are highly prevalent from 2016 to 2022. Although decreased resistance rate for both antibiotics was observed from 2016 to 2022, several resistant HierCC clusters containing hundreds of resistant isolates have shown 1.5 to 2-fold increase in frequency from 2020 to 2022 (**Supplementary Figure 5** and **Supplementary Figure 6**). Majority of the recent emerging resistant clusters carried *bla*_OXA_ genes coding for beta-lactams resistance and *tetO* gene for tetracycline resistance. These clusters have become dominant in more recent years, which would have been ignored by traditional tracking methods. The multilevel HierCC typing scheme provides a mean to differentiate and track these clusters, pinpointing emergent resistance and the genetic background of the strains from which the resistance arose. It enables precise surveillance of resistant clusters and better design of targeted prevention and control strategies.

We further identified sequential acquisition of resistance using network analysis. Subset of beta-lactam resistant clusters at HC264 acquired tetracycline resistance genes and evolved into resistance to multiple antibiotics as resistant clusters at the higher HierCC levels (**Figure 6C**). Similarly, prevalence of quinolones resistance in *C. jejuni* was around ∼21% in the USA dataset with the most prevalent resistant mechanism detected being mutations in *gyrA*. Our hierarchical network analysis found that clusters resistant to other antibiotics can acquire resistance to quinolones rapidly. Shown as an example in **Figure 6D**, two subsets of narrow-spectrum beta-lactams resistant HC264 cluster 1115 (HC55 cluster 139 and HC19 cluster 711) acquired additional resistance to quinolones. The multilevel HierCC typing scheme had provided a more targeted platform for analysing emergence of resistance and changing trends of resistance.

### Monitoring *50S_L22* mutations in response to selection pressure of macrolides

In *C. jejuni*, phenotypic macrolide resistance is mostly associated with *erm*(B) and mutations in the 23S rRNA [77–79]. *50S_L22* mutations alone do not confer phenotype macrolide resistance in *C. jejuni* [80]. Previous studies have reported the synergy between efflux pump CmeABC and *50S_L22(A103V)* mutation may lead to resistance to macrolides in *C. jejuni* [81]. However, by utilising phenotypic data from NCBI Pathogen Detection database, we have found all the isolates carrying with both *cmeABC* gene and *50S_L22(A103V)* mutation are susceptible to both azithromycin and erythromycin. Nonetheless, *50S_L22* mutations serves as a reliable marker of macrolide selection pressure. During an *in vitro* stepwise exposure experiment, when exposing *C. jejuni* strains to increasing dosages of erythromycin or tylosin, *50S_L22* mutations occurred earlier compared to 23S rRNA mutations [82]. Therefore, *50S_L22* mutations have likely arisen through selection pressure from macrolides. However, mutation of the *50S_L22* gene downregulates motility and metabolism genes, which is a significant fitness cost for mutated strains to compete with strains without the mutation under no selection pressure [83]. These findings may account for why the 2016– 2022 USA dataset contains a high proportion of *50S_L22* mutations but very few 23S rRNA mutations.

By utilising the temporal trend analysis, we identified *50S_L22* mutation carrying clusters correlated with the increase of macrolide usage at specific time frames. The increased percentage of genome carrying *50S_L22* mutations from 2016 to 2022 indicated elevated selection pressure from macrolides usage in the USA. This can be correlated with the widespread usage of macrolides in the US chicken and cattle farming industry [19,69,75]. In the USA, *50S_L22* mutations increased significantly from the 2016-17 period to 2018-2019 period, and again from the 2018-2019 period to the 2020-2022 period (**Figure 4**). These findings highlighted the importance of continuous monitoring of AMR or AMR associated mutations for better-informed agricultural legislations and public health safety. The multilevel HierCC typing scheme enable tracking clusters carrying *50S_L22* mutations.

## Conclusions

In this study, we built a multilevel HierCC typing scheme to provide flexible, stable and standardised genomic typing for *C. jejuni*. We formulated a statistical process to identify epidemiologically meaningful HierCC level for both long and short-term epidemiological analysis. The multilevel design provided flexible typing resolution while maintaining the stability of the overall nomenclature. The nine epidemiologically meaningful levels also provided a reference for comparison between single-level scheme based local studies.

The scheme has shown promising utility in identifying continent or country specific clusters and farm-animal specific clusters in *C. jejuni* populations. Importantly, the scheme enabled tracking and monitoring of AMR trends to medically important antibiotic classes. Using surveillance genome data from the USA and the UK we showcased the robustness of the scheme in AMR surveillance. We believe the multilevel HierCC typing scheme has the potential to be integrated into a continuous and comprehensive *C. jejuni* surveillance program, supporting data-driven decision making-for the prevention and control of *C. jejuni* globally.

## Data availability

The UNSW *C. jejuni* cgMLST scheme is available at the Chewie Nomenclature Serve (**Chewie-NS**) (https://chewbbaca.online). A guide to download and implement the *C. jejuni* multilevel HierCC typing scheme has been created and is freely available at GitHub (https://github.com/LanLab/C.jejuni-multilevel-HierCC-typing-scheme ).

## Author’s contributions

R.W.: collected, processed, and analysed the data. constructed the python script used in this project, writing, and editing of the manuscript. S.K.: supervised the project, writing and editing of the manuscript. M.P.: supervised the project and provided feedback on the manuscript. L.Z.: conceived the project with R.L. and provided feedback on the manuscript. R.L.: conceived and supervised the project and writing and editing of the manuscript.

## Acknowledgements

The authors would like to thank the UNSW ResTech Technology Services Team for their ongoing assistance with the high-powered computing and data management systems.

