## Supplementary material for "Multilevel HierCC typing scheme for global genomic epidemiology and antimicrobial resistance surveillance of *Campylobacter jejuni*": Supplymentary Figures

**Supplementary Figures**

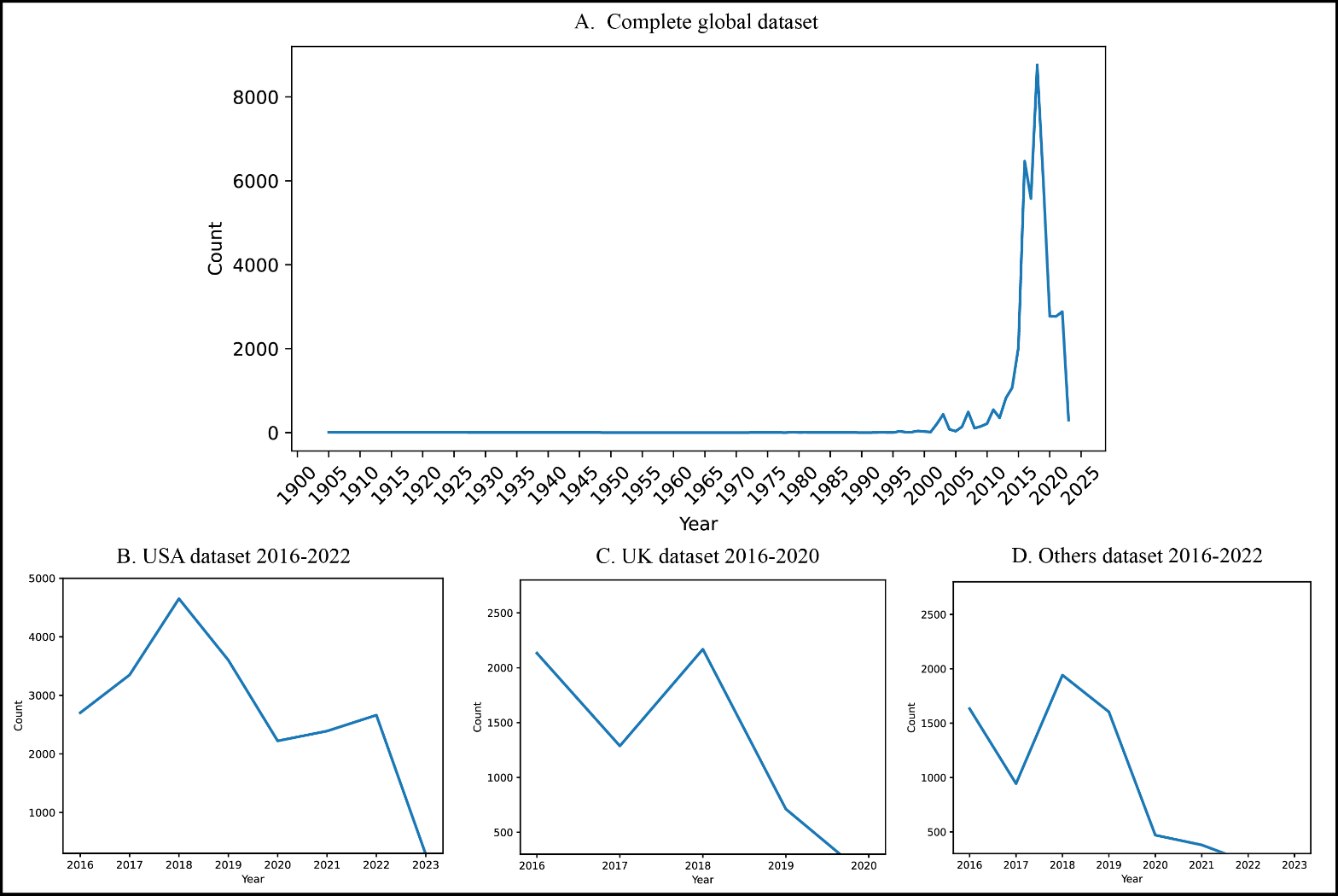

**Supplementary Figure 1 Sampling of global dataset.**

Sampling of data from the global dataset from 1905 – 2023 (A). Three subsets of data were identified based on country information: B. USA dataset from 2016-2022, C. UK dataset from 2016-2022, and D. isolates collected from other countries from 2016-2022.

**Supplementary Figure 2 ARI comparison between clusters in each HierCC level with 7MLST STs.**

Adjusted Rand Index (ADI)

| HierCC levels | Adjusted_Rand_Score |
| --- | --- |
| HC0 | 0.0037899 |
| HC1 | 0.0126562 |
| HC2 | 0.0203747 |
| HC3 | 0.0283054 |
| HC4 | 0.0363172 |
| HC5 | 0.0493332 |
| HC6 | 0.0587232 |
| HC7 | 0.0695074 |
| HC8 | 0.0816791 |
| HC9 | 0.100165 |
| HC10 | 0.111297 |
| HC11 | 0.121975 |
| HC12 | 0.129137 |
| HC13 | 0.144809 |
| HC14 | 0.153167 |
| HC15 | 0.158353 |
| HC16 | 0.166007 |
| HC17 | 0.172789 |
| HC18 | 0.17888 |
| HC19 | 0.183569 |
| HC20 | 0.190819 |
| HC21 | 0.20746 |
| HC22 | 0.215135 |
| HC23 | 0.224296 |
| HC24 | 0.234766 |
| HC25 | 0.240891 |
| HC26 | 0.246944 |
| HC27 | 0.2519 |
| HC28 | 0.257091 |
| HC29 | 0.266426 |
| HC30 | 0.273392 |
| HC31 | 0.277952 |
| HC32 | 0.281173 |
| HC33 | 0.286631 |
| HC34 | 0.289751 |
| HC35 | 0.295741 |
| HC36 | 0.297142 |
| HC37 | 0.298236 |
| HC38 | 0.309592 |
| HC39 | 0.314122 |
| HC40 | 0.31715 |
| HC41 | 0.320753 |
| HC42 | 0.323706 |
| HC43 | 0.332326 |
| HC44 | 0.335792 |
| HC45 | 0.338238 |
| HC46 | 0.34503 |
| HC47 | 0.348242 |
| HC48 | 0.351207 |
| HC49 | 0.353698 |
| HC50 | 0.356259 |
| HC51 | 0.368246 |
| HC52 | 0.374062 |
| HC53 | 0.377645 |
| HC54 | 0.380461 |
| HC55 | 0.382041 |
| HC56 | 0.385728 |
| HC57 | 0.389314 |
| HC58 | 0.391222 |
| HC59 | 0.427909 |
| HC60 | 0.428962 |
| HC61 | 0.433575 |
| HC62 | 0.434809 |
| HC63 | 0.438465 |
| HC64 | 0.446401 |
| HC65 | 0.45045 |
| HC66 | 0.453257 |
| HC67 | 0.455019 |
| HC68 | 0.45795 |
| HC69 | 0.458935 |
| HC70 | 0.461272 |
| HC71 | 0.463307 |
| HC72 | 0.469817 |
| HC73 | 0.473496 |
| HC74 | 0.473943 |
| HC75 | 0.473677 |
| HC76 | 0.474639 |
| HC77 | 0.475215 |
| HC78 | 0.475842 |
| HC79 | 0.477659 |
| HC80 | 0.477474 |
| HC81 | 0.477684 |
| HC82 | 0.480846 |
| HC83 | 0.472363 |
| HC84 | 0.476827 |
| HC85 | 0.477154 |
| HC86 | 0.478936 |
| HC87 | 0.480174 |
| HC88 | 0.482809 |
| HC89 | 0.480082 |
| HC90 | 0.48005 |
| HC91 | 0.48293 |
| HC92 | 0.483445 |
| HC93 | 0.485772 |
| HC94 | 0.49099 |
| HC95 | 0.491638 |
| HC96 | 0.492412 |
| HC97 | 0.493086 |
| HC98 | 0.493493 |
| HC99 | 0.4953 |
| HC100 | 0.495452 |
| HC101 | 0.495675 |
| HC102 | 0.496698 |
| HC103 | 0.496825 |
| HC104 | 0.508275 |
| HC105 | 0.508877 |
| HC106 | 0.508724 |
| HC107 | 0.502096 |
| HC108 | 0.502466 |
| HC109 | 0.502892 |
| HC110 | 0.503315 |
| HC111 | 0.50311 |
| HC112 | 0.50352 |
| HC113 | 0.50857 |
| HC114 | 0.509623 |
| HC115 | 0.510308 |
| HC116 | 0.511262 |
| HC117 | 0.511535 |
| HC118 | 0.512093 |
| HC119 | 0.512259 |
| HC120 | 0.514222 |
| HC121 | 0.514118 |
| HC122 | 0.513565 |
| HC123 | 0.513712 |
| HC124 | 0.559613 |
| HC125 | 0.559808 |
| HC126 | 0.560096 |
| HC127 | 0.563484 |
| HC128 | 0.563475 |
| HC129 | 0.563722 |
| HC130 | 0.564073 |
| HC131 | 0.563786 |
| HC132 | 0.56418 |
| HC133 | 0.564447 |
| HC134 | 0.564533 |
| HC135 | 0.562818 |
| HC136 | 0.562306 |
| HC137 | 0.579711 |
| HC138 | 0.5797 |
| HC139 | 0.580205 |
| HC140 | 0.579312 |
| HC141 | 0.579557 |
| HC142 | 0.59877 |
| HC143 | 0.598993 |
| HC144 | 0.598983 |
| HC145 | 0.599118 |
| HC146 | 0.600448 |
| HC147 | 0.603207 |
| HC148 | 0.60289 |
| HC149 | 0.605517 |
| HC150 | 0.60554 |
| HC151 | 0.605542 |
| HC152 | 0.605625 |
| HC153 | 0.605712 |
| HC154 | 0.610881 |
| HC155 | 0.610038 |
| HC156 | 0.615385 |
| HC157 | 0.618524 |
| HC158 | 0.618289 |
| HC159 | 0.618168 |
| HC160 | 0.617618 |
| HC161 | 0.617833 |
| HC162 | 0.619585 |
| HC163 | 0.620893 |
| HC164 | 0.620903 |
| HC165 | 0.624725 |
| HC166 | 0.624506 |
| HC167 | 0.624473 |
| HC168 | 0.625029 |
| HC169 | 0.624963 |
| HC170 | 0.62514 |
| HC171 | 0.625116 |
| HC172 | 0.626408 |
| HC173 | 0.62715 |
| HC174 | 0.627599 |
| HC175 | 0.627612 |
| HC176 | 0.626676 |
| HC177 | 0.626605 |
| HC178 | 0.626859 |
| HC179 | 0.626838 |
| HC180 | 0.626918 |
| HC181 | 0.626808 |
| HC182 | 0.626774 |
| HC183 | 0.627838 |
| HC184 | 0.627817 |
| HC185 | 0.625628 |
| HC186 | 0.603238 |
| HC187 | 0.603028 |
| HC188 | 0.603113 |
| HC189 | 0.603092 |
| HC190 | 0.610707 |
| HC191 | 0.610728 |
| HC192 | 0.610679 |
| HC193 | 0.610936 |
| HC194 | 0.611006 |
| HC195 | 0.612029 |
| HC196 | 0.612287 |
| HC197 | 0.612313 |
| HC198 | 0.612183 |
| HC199 | 0.611792 |

The black line indicates the adjusted rand index (ARI) between the clusters at each HierCC level (0 - 1160) and the 7MLST ST of the global dataset (n=63102 genomes). ARI was above 0.5 between HC 104 to HC 335 and peaked at HC 264 at 0.66. All ARI between 7MLST and HierCC level were above 0.

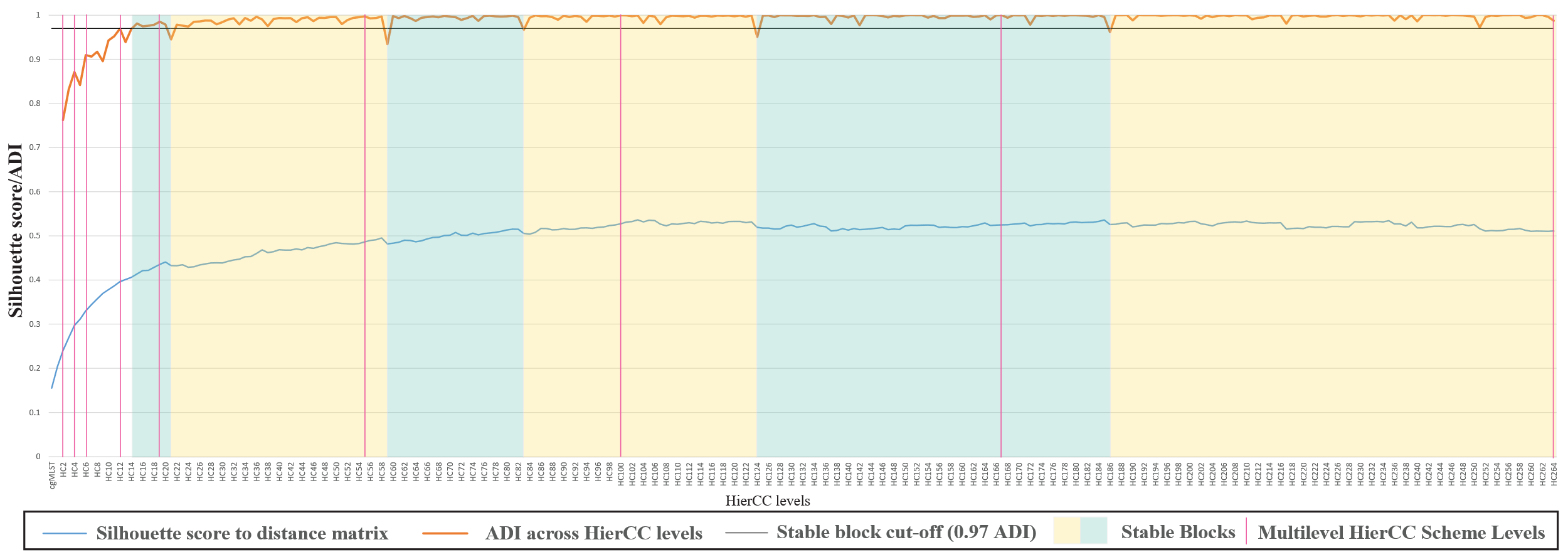

**Supplementary Figure 3 Identification of HierCC levels that best dividing the population structure with different resolutions for epidemiological typing.**

ARI was measured for every two HierCC levels from to HC0 to HC264. A soft cut-off of 0.97 ARI has been set to identify HierCC levels with unstable dips in ARI. HierCC levels between these dips are considered as a part of a “stable block”. HierCC levels with the highest Silhouette score with the cgMLST allelic distance-matrix within each stable block were selected as epidemiologically meaningful levels. The location of the selected levels (HC167, HC100, HC55, HC19, HC12, HC6, HC4, and HC2) are indicated with a red vertical line

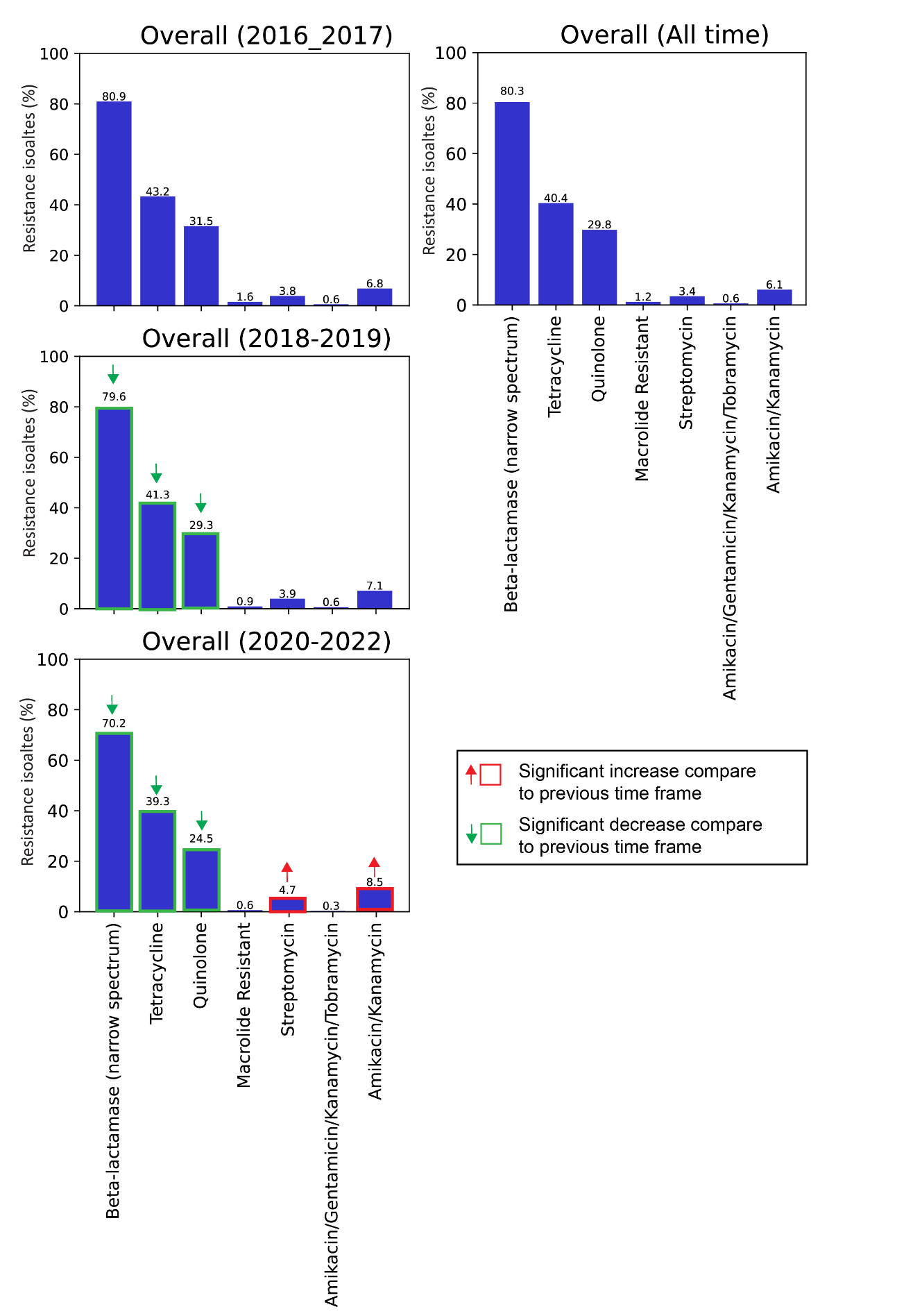
Percentage of resistance isolates to seven antibiotic classes or combinations (narrow spectrum beta-lactams, tetracyclines, quinolones, macrolides, streptomycin, amikacin/gentamycin/kanamycin/tobramycin, amikacin/kanamycin) within the global dataset. Resistance to narrow spectrum beta-lactams, tetracyclines and quinolones has decreased significantly from 2016 to 2022. Resistance to streptomycin and the amikacin/kanamycin combination has increased significantly. The resistance to a given antibiotic class in different time frames was compared using the binomial test. Significance was assessed with the Bonferroni corrected *p* < 0.0008. Significant increase or decrease in resistance is indicated by an upwards red arrow or a downwards green arrow, respectively.

**Supplementary Figure 4 Resistant isolates to seven antibiotic class or combinations in the global dataset in three timeframes.**

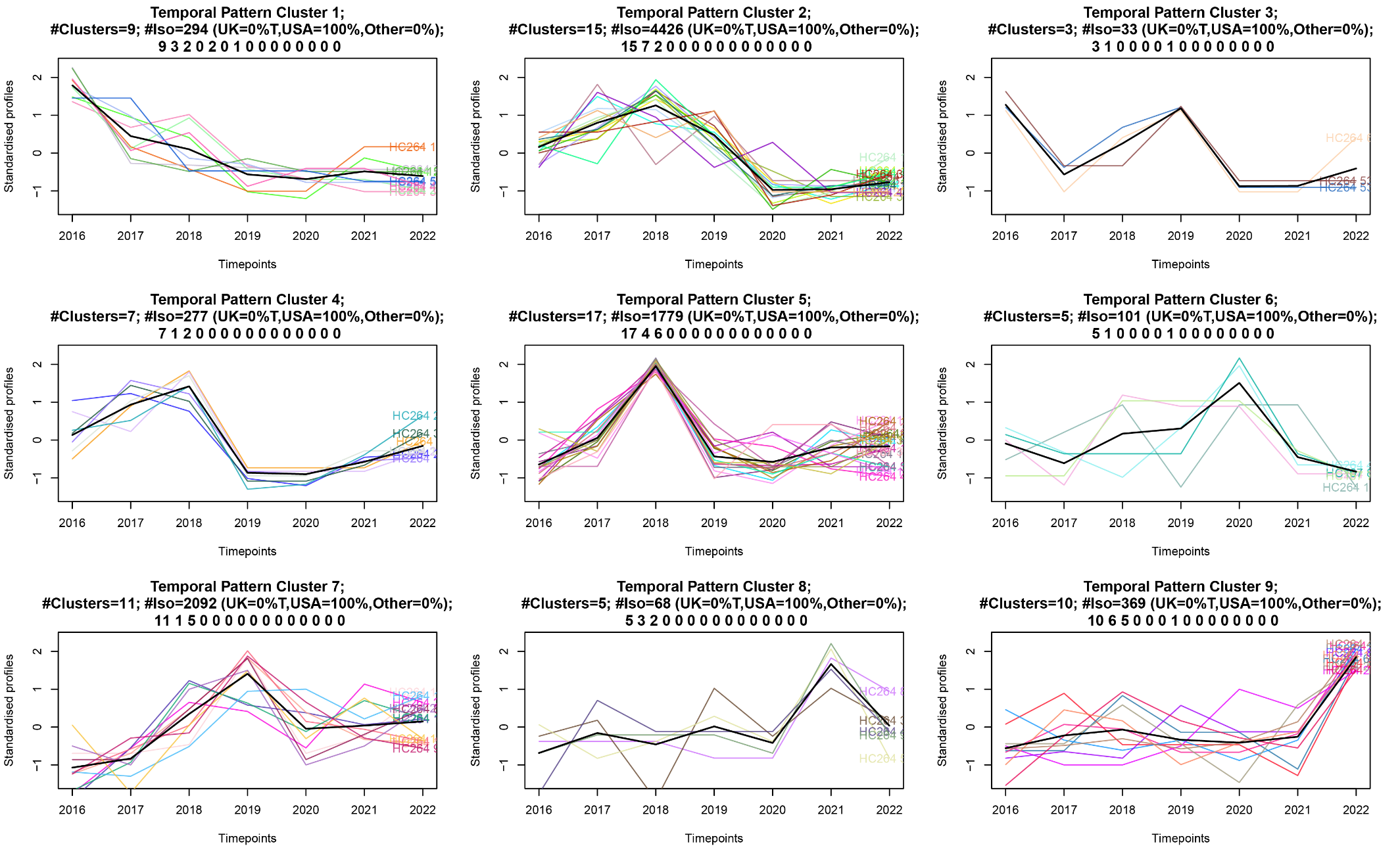

**Supplementary Figure 5 Temporal resistant pattern of beta-lactams in the USA in 2016-2022.**

Temporal trends of HierCC clusters resistant to narrow-spectrum beta-lactams between 2016-2022 in the USA dataset. Nine TPC contained 82 narrow-spectrum beta-lactams resistant clusters. An individual graph with separate header was created for each TPC. Each header indicates the number of resistant clusters (#clusters) within the TPC, total number of isolates (#Iso) and the country of origin of the isolates (UK, USA or other). The series of numbers at the bottom of the header indicated the number of HierCC clusters resistant to a particular antibiotic class or combination. (e.g.: 9 3 2 0 2 0 1 0 0 0 0 0 0 0 0 in the first graph, here 9 means all nine clusters in this TPC are resistant to narrow-spectrum beta-lactamase). The order of the 15 antibiotic classes or combinations are as follows: 1. Beta-lactamase (narrow-spectrum), 2. Tetracycline, 3. Quinolone, 4. Macrolide, 5. Streptomycin, 6. Amikacin/Gentamicin/Kanamycin/Tobramycin, 7. Amikacin/Kanamycin, 8. Kanamycin/Tobramycin, 9. Amikacin/Kanamycin/Tobramycin, 10. Lincosamides, 11. Lincosamide/Streptogramin, 12. Chloramphenicol, 13. Chloramphenicol/Florfenicol, 14. Sulfonamide, 15. Florfenicol/Oxazolidinone.

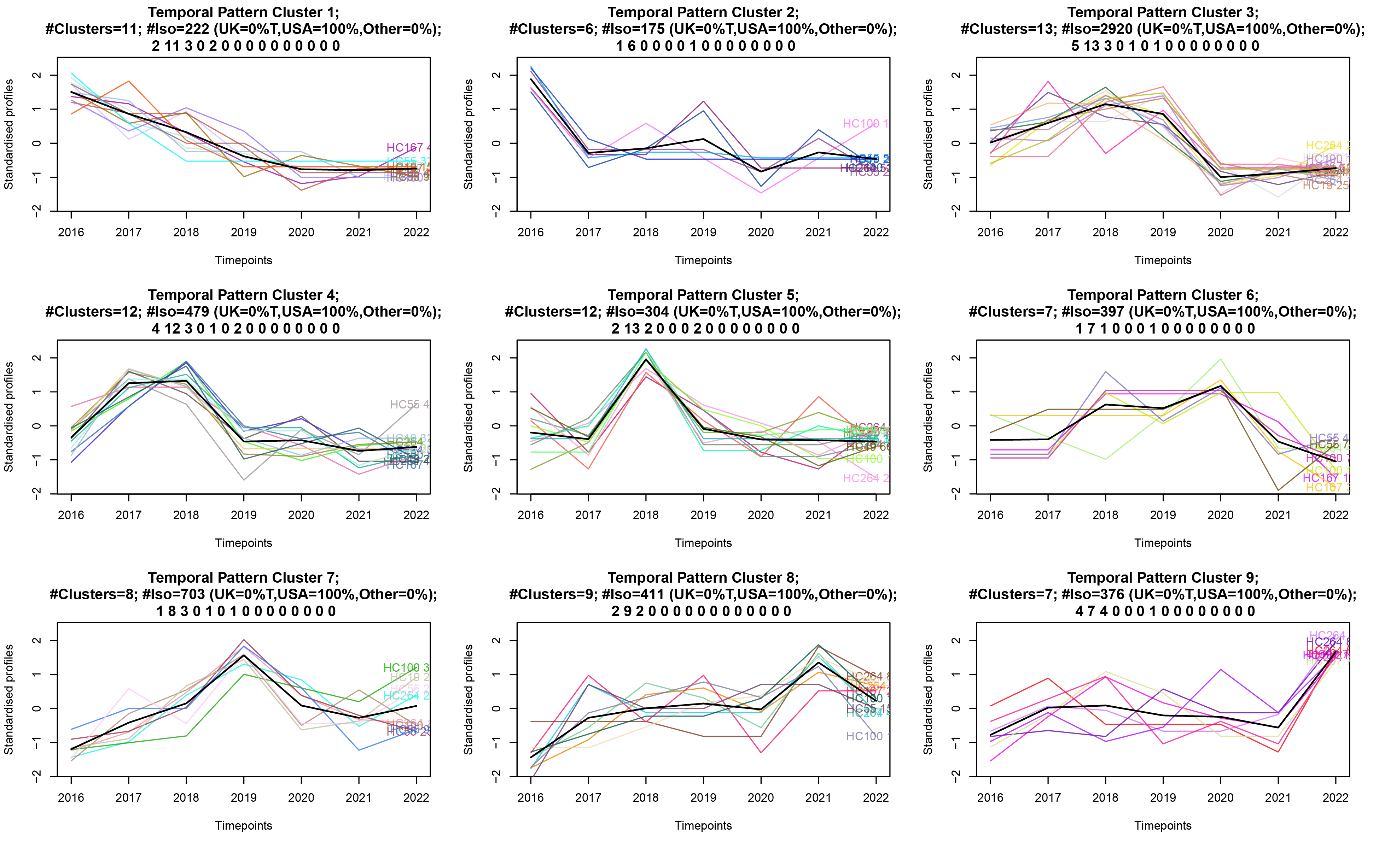
Temporal trends of HierCC clusters resistant to tetracycline between 2016-2022 in the USA dataset. Nine TPC contained 85 tetracycline resistant clusters. For each TPC, temporal profile of each unique resistant HierCC cluster is the standardised count of isolates per year, which was represented by a unique coloured line. The weighted average temporal pattern of all resistant clusters in a TPC is represented by a bold-black line. An individual graph with separate header was created for each TPC. Each header indicates the number of resistant clusters (#clusters) within the TPC, total number of isolates (#Iso) and the country of origin of the isolates (UK, USA or other). The series of numbers at the bottom of the header indicated the number of HierCC clusters resistant to a particular antibiotic class or combination. (e.g.: 2 11 3 0 2 0 0 0 0 0 0 0 0 0 0 in the first graph, here 2 means 2 of the 11 clusters in this TPC are resistant to narrow-spectrum beta-lactamase). The order of the 15 antibiotic classes or combinations are as follows: 1. Beta-lactamase (narrow-spectrum), 2. Tetracycline, 3. Quinolone, 4. Macrolide, 5. Streptomycin, 6. Amikacin/Gentamicin/Kanamycin/Tobramycin, 7. Amikacin/Kanamycin, 8. Kanamycin/Tobramycin, 9. Amikacin/Kanamycin/Tobramycin, 10. Lincosamides, 11. Lincosamide/Streptogramin, 12. Chloramphenicol, 13. Chloramphenicol/Florfenicol, 14. Sulfonamide, 15. Florfenicol/Oxazolidinone.

**Supplementary Figure 6 Temporal resistant pattern of tetracyclines in the USA in 2016-2022.**

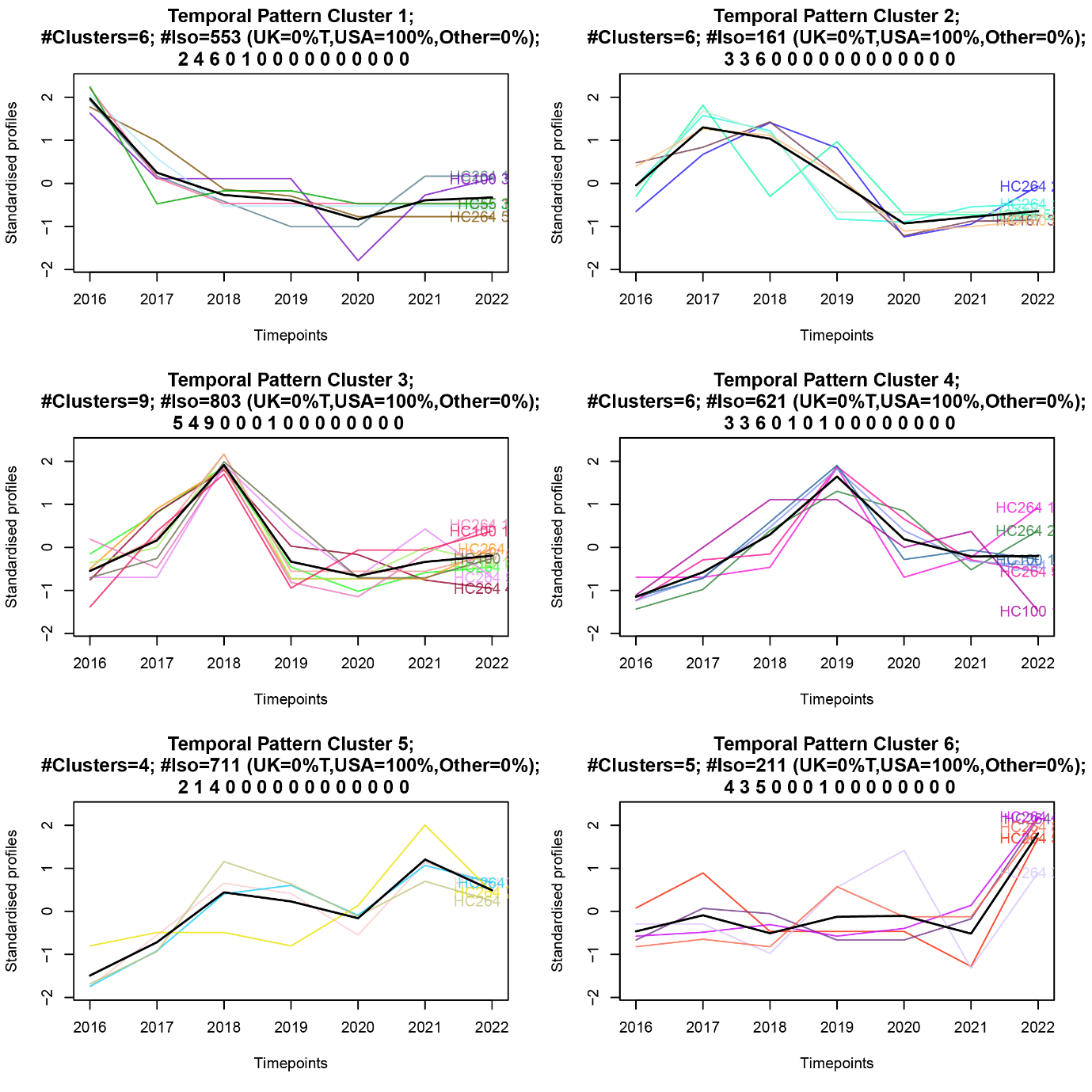

**Supplementary Figure 7 Temporal resistant pattern of quinolones in the USA in 2016-2022***.*

Temporal trends of HierCC clusters resistant to quinolones between 2016-2022 in the USA dataset. Six TPC contained 36 quinolones resistant clusters. For each TPC, temporal profile of each unique resistant HierCC cluster is the standardised count of isolates per year, which was represented by a unique coloured line. The weighted average temporal pattern of all resistant clusters in a TPC is represented by a bold-black line. An individual graph with separate header was created for each TPC. Each header indicates the number of resistant clusters (#clusters) within the TPC, total number of isolates (#Iso) and the country of origin of the isolates (UK, USA or other). The series of numbers at the bottom of the header indicated the number of HierCC clusters resistant to a particular antibiotic class or combination. (e.g.: 2 4 6 0 1 0 6 0 0 0 0 0 0 0 0 in the first graph, here 2 means 2 of the 6 clusters in this TPC are resistant to narrow-spectrum beta-lactamase). The order of the 15 antibiotic classes or combinations are as follows: 1. Beta-lactamase (narrow-spectrum), 2. Tetracycline, 3. Quinolone, 4. Macrolide, 5. Streptomycin, 6. Amikacin/Gentamicin/Kanamycin/Tobramycin, 7. Amikacin/Kanamycin, 8. Kanamycin/Tobramycin, 9. Amikacin/Kanamycin/Tobramycin, 10. Lincosamides, 11. Lincosamide/Streptogramin, 12. Chloramphenicol, 13. Chloramphenicol/Florfenicol, 14. Sulfonamide, 15. Florfenicol/Oxazolidinone. The optimal number of TPCs was determined through a combination of the silhouette index and manual inspection of the temporal delineation of the clusters, TPCs were then generated using the unsupervised c-means clustering algorithm.
