## Supplementary material for "Multilevel HierCC typing scheme for global genomic epidemiology and antimicrobial resistance surveillance of *Campylobacter jejuni*": Supplymentary Methods

**Supplementary Methods**

**1.1 Core loci validation for the global *C. jejuni* cgMLST**

The assembled genomes were filtered based on the following quality thresholds: genomes size between 1509232 bp and 1858564 bp, a maximum of 119 contigs, a maximum of 292243 bp for N50, and a maximum L50 of 16.5 contigs.

**1.2 Core loci validation for the global *C. jejuni* cgMLST**

We downloaded 346 publicly available *C. jejuni* complete whole genome sequences from NCBI (<https://www.ncbi.nlm.nih.gov/>) and a training file generated by Prodigal v2.6.3 from the *C. jejuni* reference genome (RefSeq Accession GCF_000304375.1) to establish the initial schema seeds for the global *C. jejuni* global cgMLST scheme. The Comprehensive and Highly Efficient Workflow BSR-Based Allele Calling Algorithm (chewBBACA v3.3.2) predicted 3920 coding sequences (CDS) from the 346 complete genomes. Allele calling was performed with a Blast Score Ratio of 0.6 to redetermine the core loci from the schema seeds. For a CDS to be considered a core locus, it must be present in >=95% of the diverse representative dataset (n=2587).

**1.3 Order of the antibiotics and antibiotic classes in resistance score**

1. beta-lactamase (narrow-spectrum), 2. tetracycline, 3. quinolone, 4. macrolide, 5. streptomycin, 6. amikacin/gentamicin/kanamycin/tobramycin, 7. amikacin/kanamycin, 8. kanamycin/tobramycin, 9. amikacin/kanamycin/tobramycin, 10. lincosamides, 11. lincosamide/streptogramin, 12. chloramphenicol, 13. chloramphenicol/florfenicol, 14. sulfonamide, 15. florfenicol/oxazolidinone.
