## Supplementary material for "Multilevel HierCC typing scheme for global genomic epidemiology and antimicrobial resistance surveillance of *Campylobacter jejuni*": Supplymentary Tables

**Supplementary Tables**

**Supplementary table 2 Metrics of continent-specific clusters at different HierCC levels**

| HierCC level | Percentage of isolates in continent-specific clusters | Number of continent-specific clusters | Number of isolates in continent specific clusters | Number of isolates in major clusters (>=10 isolates) |
| --- | --- | --- | --- | --- |
| HC2 | 93.39% | 308 | 8501 | 9103 |
| HC4 | 95.22% | 466 | 14371 | 15093 |
| HC6 | 94.71% | 592 | 19392 | 20476 |
| HC12 | 95.85% | 731 | 29135 | 30395 |
| HC19 | 93.38% | 749 | 34549 | 36998 |
| HC55 | 88.02% | 593 | 44012 | 50002 |
| HC100 | 83.95% | 425 | 46022 | 54823 |
| HC167 | 63.17% | 302 | 36523 | 57817 |
| HC264 | 45.46% | 182 | 27173 | 59779 |

**Supplementary table 3 Metrics of Host-specific clusters at different HierCC levels**

| HierCC level | Percentage of isolates in farm-animal-associated clusters | Number of farm-animal-associated clusters | Number of isolates in farm-animal-associated clusters | Number of isolates in >=10 isolates clusters |
| --- | --- | --- | --- | --- |
| HC2 | 0.8914 | 62 | 1247 | 1399 |
| HC4 | 0.8356 | 103 | 2449 | 2931 |
| HC6 | 0.7644 | 137 | 3351 | 4384 |
| HC12 | 0.7192 | 197 | 5270 | 7328 |
| HC19 | 0.7062 | 239 | 6952 | 9844 |
| HC55 | 0.7052 | 259 | 11307 | 16034 |
| HC100 | 0.6975 | 208 | 12761 | 18296 |
| HC167 | 0.6207 | 150 | 12299 | 19814 |
| HC264 | 0.5405 | 94 | 11252 | 20817 |

**Supplementary table 4 Accuracy of AbritAMR phenotype interpretation against *C. jejuni***

| Antibiotic | Phenotype Resistance | AbritAMR interpretation of resistance | Accuracy |
| --- | --- | --- | --- |
| ampicillin | 21 | 21 | 100.00% |
| ciprofloxacin | 658 | 653 | 99.24% |
| tetracyclines | 1528 | 1509 | 98.76% |
| azithromycin | 75 | 74 | 98.67% |
| nalidixic acid | 634 | 624 | 98.42% |
| erythromycin | 78 | 76 | 97.44% |
| gentamicin | 29 | 28 | 96.55% |
| telithromycin | 58 | 54 | 93.10% |
| **Average** |  |  | **97.77%** |

| Drug class | UK 2016-2019 | USA 2016-2022 | Others 2016-2022 |
| --- | --- | --- | --- |
| Narrow-spectrum beta-lactams | *bla*_OXA-193_ (60.12%), *bla*_OXA-461_ (7.66%), *bla*_OXA-184_ (7.62%), bla_OXA-61_ (6.80%) | *bla*_OXA-193_ (58.54%), *bla*_OXA-184_ (8.37%), *bla*_OXA-61_ (6.91%) | *bla*_OXA-193_ (59.52%), *bla*_OXA-461_ (5.94%), *bla*_OXA-61_ (5.03%) |
| Tetracyclines | *tetO* (99.37%) | *tetO* (99.67%) | *tetO* (97.86%) |
| Quinolones | *gyrA*_T86I (97.49%) | *gyrA*_T86I (98.58%) | *gyrA*_T86I (94.29%) |
| Macrolides | *23S*_A2075G (88.89%), *23S*_A2074T (3.70%) | *23S*_A2075G 90.99%), *23S*_A2074T (6.01%) | *23S*_A2074T (42.86%), *23S*_A2075G (35.71%) |

**Supplementary table 5 Most common resistance determinants for UK, USA and other countries**

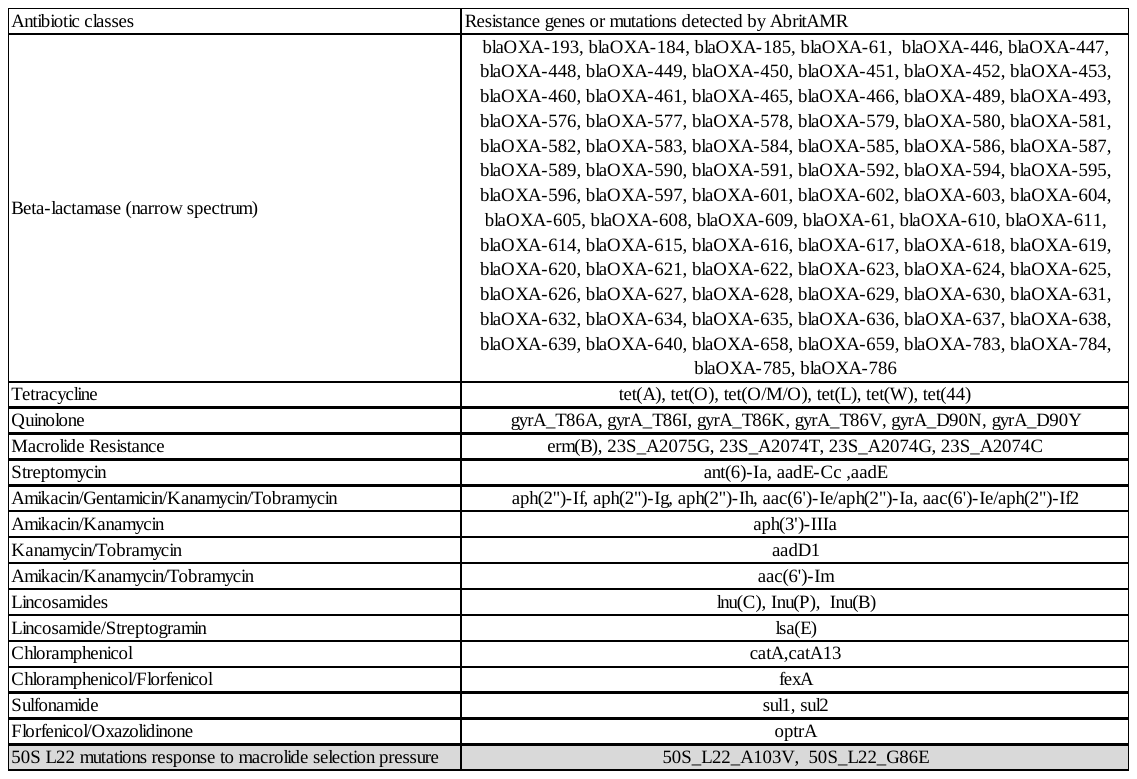

**Supplementary Table 6 Resistance genes and mutations detected by AbritAMR in the global dataset**

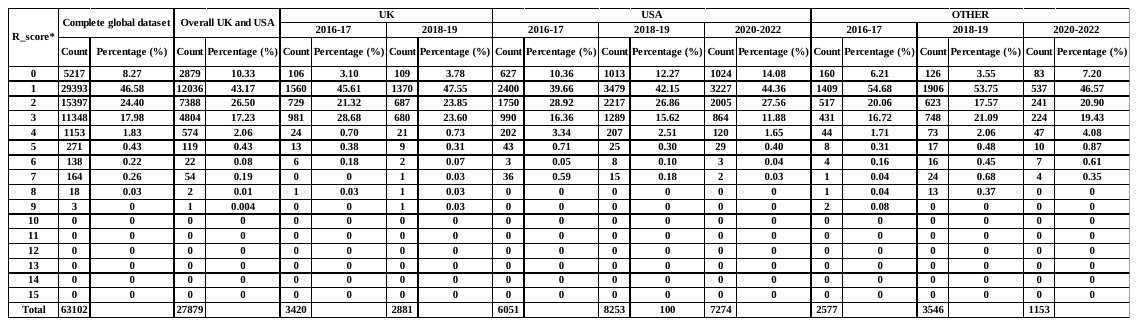
*R_score: resistance score indicating the number of antibiotics to which an isolate was predicted to be resistant.

**Supplementary Table 7 Resistance score of the global dataset, the UK, the USA, and other countries from 2016-2022**

**Supplementary table 8 Resistant clusters to medically important antibiotics in the USA from 2016-2022.**

| Drug | HC264 | HC167 | HC100 | HC55 | HC19 | HC12 | HC6 | HC4 | HC2 | 2016-2022 resistant clusters |
| --- | --- | --- | --- | --- | --- | --- | --- | --- | --- | --- |
| Beta-lactams (narrow-spectrum) | 78 | 2 | 0 | 1 | 0 | 0 | 0 | 0 | 0 | 81 |
| Tetracyclines | 27 | 11 | 12 | 15 | 19 | 0 | 2 | 0 | 0 | 86 |
| Quinolones | 23 | 1 | 2 | 4 | 5 | 1 | 0 | 0 | 0 | 36 |
| Macrolide | 1 | 0 | 1 | 0 | 1 | 0 | 0 | 0 | 0 | 3 |
| Total |  |  |  |  |  |  |  |  |  | **206** |
